# Zebrafish screen of high-confidence effector genes at insomnia GWAS loci implicates conserved regulators of sleep-wake behaviors

**DOI:** 10.1101/2022.10.05.511011

**Authors:** Amber J. Zimmerman, Fusun Doldur-Balli, Brendan T. Keenan, Zoe Y. Shetty, Justin Palermo, Alessandra Chesi, Shilpa Sonti, Matthew C. Pahl, Elizabeth B. Brown, James A. Pippin, Andrew D. Wells, Olivia J. Veatch, Diego R. Mazzotti, Anitra Krishnan, Phillip R. Gehrman, Alex C. Keene, Struan F.A. Grant, Allan I. Pack

## Abstract

Recent large-scale human genome-wide association studies (GWAS) for insomnia have identified more than 200 significant loci. The functional relevance of these loci to the pathogenesis of insomnia is largely unknown. GWAS signals are typically non-coding variants, which are often arbitrarily annotated to the nearest protein-coding gene; however, due to 3D chromatin structure, variants can interact with more distal genes driving their function. The distal gene may, therefore, represent the true causal gene influencing the phenotype. By integrating our high-resolution chromatin interaction maps from neural progenitor cells with phenotypic data from a *Drosophila* RNAi screen, we prioritized candidate genes that we hypothesized would have deep phylogenetic conservation of sleep function. To determine the conservation of these candidate genes in the context of vertebrate sleep and their relevance to insomnia-like behaviors, we performed CRISPR-Cas9 mutagenesis in larval zebrafish for six highly conserved candidate genes and examined sleep-wake behaviors using automated video-tracking. CRISPR mutation of zebrafish orthologs of *MEIS1* and *SKIV2L* produced insomnia-like behaviors, while mutation of *ARFGAP2* impaired activity and development in our larval zebrafish model, demonstrating the importance of performing functional validation of GWAS-implicated effector genes to reveal genes influencing disease-relevant mechanisms.

## 1. Introduction

Chronic sleep disruption is linked to a variety of negative health sequelae, including impaired metabolic and cognitive function. Nearly one-third of the adult population reports chronic sleep disturbance and symptoms of insomnia (Stranges *et al*., 2012). Insomnia is characterized as a combination of difficulty initiating sleep (increased sleep latency), and/or difficulty maintaining sleep accompanied by daytime consequences (e.g. fatigue, irritability) despite ample opportunity for sleep (Association, 2013; Morin *et al*., 2015). Insomnia, along with other sleep traits including sleep duration, napping, and daytime dozing, are heritable and extremely polygenic (Dashti *et al*., 2019; Lane *et al*., 2017). Large-scale genome-wide association studies (GWAS) have revealed several hundred genomic loci for insomnia and other sleep traits (Dashti *et al*., 2019; Hammerschlag *et al*., 2017; Jansen *et al*., 2019; Lane *et al*., 2017; Watanabe *et al*., 2022). Publicly available datasets for chromatin accessibility and gene expression have aided gene mapping at these GWAS loci; however, the majority of loci are still typically positionally mapped to the nearest gene or mapped using *in silico* prediction on aggregate data (Watanabe *et al*., 2022). These approaches can misidentify the true causal effector gene(s) at a GWAS locus (Claussnitzer *et al*., 2020; Forgetta *et al*., 2022; Fulco *et al*., 2019; Lappalainen and MacArthur, 2021; Smemo *et al*., 2014; Tam *et al*., 2019), which in turn can lead to mischaracterization of mechanisms underlying insomnia and limit the utility of human genomics data for informing clinical care. Detailed fine-mapping of genome-wide significant loci should ideally be carried out in a specific cell-type setting yielding disease-relevant information to prioritize candidate effector genes (Chesi *et al*., 2019; Lasconi *et al*., 2021; Lasconi *et al*., 2022; Pahl *et al*., 2021; Su *et al*., 2020), and should be subsequently validated through functional phenotyping in model organisms to identify those with the greatest impact on disease pathogenesis.

Given the high conservation of sleep-wake behaviors, across species, model organisms can be leveraged to study these behaviors by assessing changes to sleep characteristics, which provide insight into the development of insomnia-like behaviors. High-throughput phenotyping in model organisms can greatly speed up gene prioritization for drug discovery and therapeutic development (Freeman *et al*., 2013; Hendricks *et al*., 2000; Tran and Prober, 2022). Zebrafish are an established vertebrate model organism to deploy efficient CRISPR/Cas9 mutagenesis paired with large-scale sleep phenotyping given both their genetic tractability and rapid developmental timeline (Tran and Prober, 2022). Unlike some model organisms, including mice, zebrafish sleep is diurnally regulated and primarily consolidated to the night similar to humans. Additionally, zebrafish sleep is circadian-regulated, reversable, and has a heightened arousal threshold, making them an appropriate model for assaying behavior relevant to sleep dysfunction (Barlow and Rihel, 2017; Chiu and Prober, 2013; Rihel *et al*., 2010; Tran and Prober, 2022). Moreover, genetic conservation between zebrafish and human is relatively high (Howe *et al*., 2013), as they are both vertebrates, and indeed many of the genes identified to date that regulate sleep are highly conserved (Jansen *et al*., 2019).

To precisely identify causal effector genes associated with human insomnia GWAS loci, we have developed a high-resolution method for 3D genomic mapping of GWAS variants using a disease-relevant cell type (Palermo *et al*., 2021; Su *et al*., 2021). This approach integrates RNA-seq, assay for transposase-accessible chromatin using sequencing (ATAC-seq) data, and high-resolution promoter-focused Capture C in order to identify physical contacts between putatively causal variants and open promoters at candidate effector genes (Chesi *et al*., 2019; Pahl *et al*., 2021; Su *et al*., 2020) in neural progenitor cells (NPCs) (Palermo *et al*., 2021). The identified effector genes serve as strong candidates for functional studies in model organisms.

A high-throughput screen using RNA interference (RNAi) in *Drosophila* was then used to identify effector genes that produced a significant alteration in sleep duration (Palermo *et al*., 2021). These studies in *Drosophila* produced a refined candidate gene list identifying highly conserved regulators of sleep function that are relevant to human insomnia, including *SKIV2L, GNB3, CBX1, MEIS1, TCF12 and ARFGAP2*. Of these genes, only *MEIS1* has been functionally connected to a behavioral phenotype reminiscent of insomnia (Hammerschlag *et al*., 2017; Lane *et al*., 2017; Thireau *et al*., 2017).

The current study applied CRISPR-Cas9 mutagenesis in a vertebrate model (zebrafish) (Kroll *et al*., 2021) to determine if the function of these implicated genes is conserved in vertebrates and relevant to sleep dysfunction observed in human insomnia. Since assessment of sleep in zebrafish is dependent on assessment of movement, we first determined if there was any evidence of movement abnormalities before examining sleep and then proceeded to examine sleep characteristics in CRISPR mutants.

## 2. Results

### 2.1 Identification of high-confidence insomnia effector genes

To examine the role and evolutionary conservation of genes regulating sleep, we integrated 3D genomics data (Palermo *et al*., 2021; Su *et al*., 2021) with phenotypic data from a high-throughput *Drosophila RNAi* screen(Palermo *et al*., 2021). Effector genes were defined as those with promoters residing in open chromatin regions, which also display high resolution chromatin contacts with putative insomnia causal variants associated with significant GWAS loci (**Fig. 1A**) in neural progenitor cells (NPCs) (Palermo *et al*., 2021). These genes are highly expressed in NPCs and demonstrate high conservation across human, zebrafish, and *Drosophila (Hu et al*., *2011)* (**Supplementary Table 1**), making them high-priority candidates for functional assessment.

**Fig. 1.**
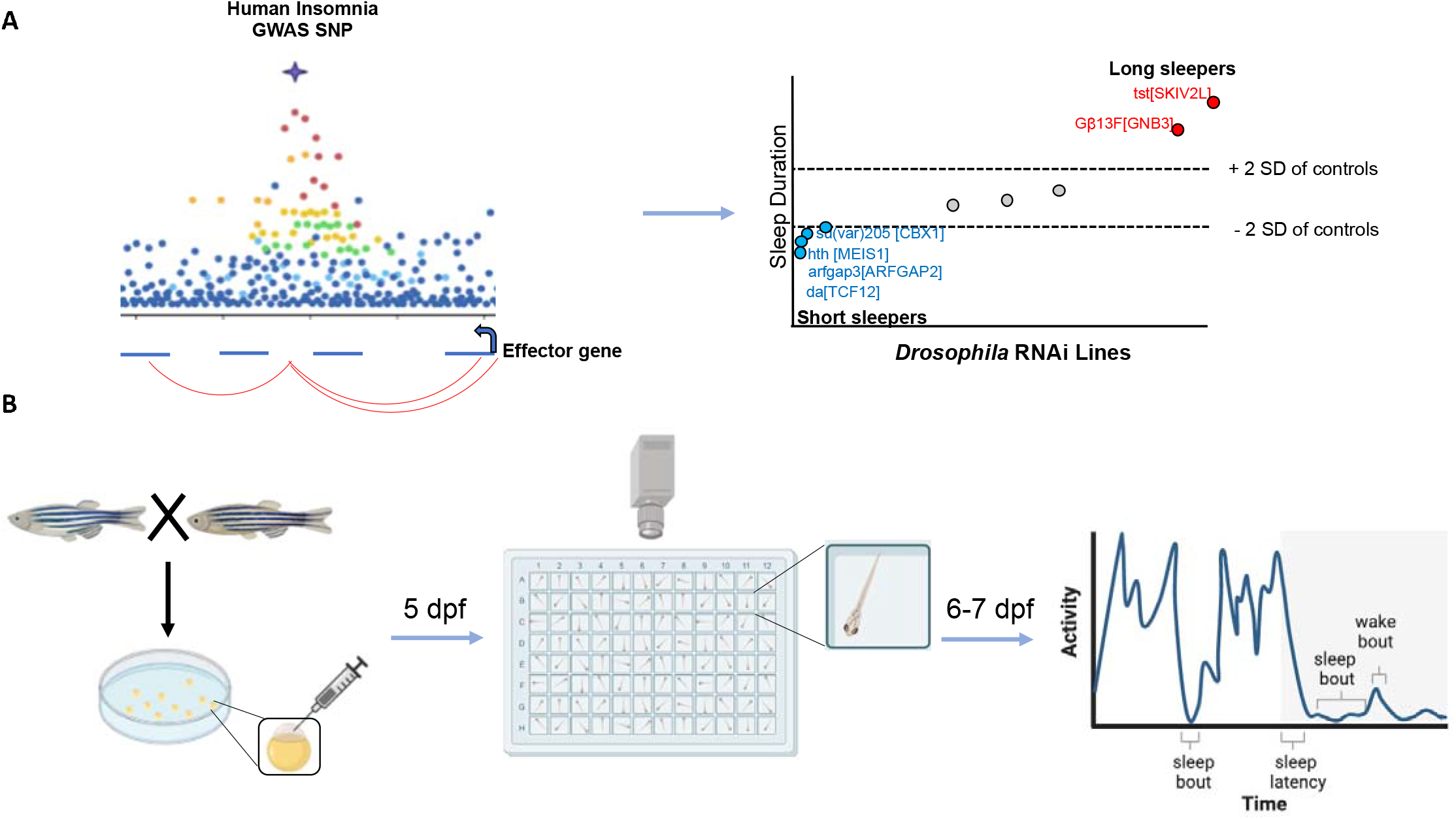
Identification of insomnia candidate genes from GWAS signals. **A**. To identify putative causal genes, insomnia GWAS variants were mapped to target effector gene in neural progenitor cells using 3D genomics, then candidate effector genes were screened using *Drosophila* RNA interference to identify those producing significant sleep phenotypes (adapted from Palermo et. al.). **B**. Single cell embryos were injected with preformed CRISPR-Cas9 complexes to create mutations in candidate effector genes. F0 larvae were tested for sleep characteristics beginning on 5 days post fertilization (dpf) and phenotyped for sleep-wake behaviors from day 6-7 using automated video tracking. Figure created in BioRender.

Our previous work performed high-throughput screening of candidate effector genes using *Drosophila* RNAi, revealing a subset of genes producing exceptionally strong sleep phenotypes (Palermo *et al*., 2021) (**Fig. 1A, right panel**), of which we chose to perform functional follow-up using zebrafish to identify conservation of function within a vertebrate model organism. The human orthologs of these genes were *SKIV2L, GNB3, CBX1, MEIS1, TCF12*, and *ARFGAP2*, which are involved in a variety of conserved cellular processes in humans involving transcriptional regulation, cellular trafficking, and signal transduction.

To determine whether these candidate effector genes exhibit strong evolutionary conservation of function related to sleep, we employed CRISPR-Cas9 mutagenesis in single-cell embryos followed by rapid behavioral screening of F0 larval zebrafish (Kroll *et al*., 2021) 5 to 7 days post fertilization (**Fig. 1B**).

### 2.2 Screening for gross movement phenotypes reveals ARFGAP2 ortholog as a neurodevelopmental gene

Given that our quantification of sleep in zebrafish is dependent on activity patterns, we first sought to examine gross motor behaviors and activity patterns to eliminate potential confounds that could contribute to the measured sleep behaviors. To do this, we measured waking activity, which is calculated as the duration of movement only during “awake” minutes (awake threshold >0.5 s/min) and serves as a proxy for general movement disruptions that may indicate gross motor changes following genetic manipulation. Since many sleep-wake behaviors are not normally distributed, we used a Wilcoxon rank sum test to test for significance between each CRISPR mutant and its own control correcting for multiple comparisons across all eleven measured sleep traits using a Hochberg step-up procedure (Hochberg, 1988; Huang and Hsu, 2007) (see **Methods**), which performs well when outcomes are correlated, as sleep traits generally are. While, s*kiv2l, gnb3a, cbx1b, meis1b, and tcf12* mutants showed no significant (*P* > 0.05 following multiple comparisons) changes to daytime or nighttime waking activity (**Fig. 2A-E, G-K**), *Gnb3a* (**Fig. 2B**), *cbx1b* (**Fig. 2C**), and *tcf12* (**Fig. 2E**) mutants showed mildly increased nighttime waking activity that did not reach the threshold for significance following multiple comparisons, which may represent a subtle hyperactive phenotype during the night. Although we observed no difference in nighttime waking activity of *arfgap2* mutants (mean difference = -0.03 s/awake minute (−0.12, 0.06, *P* = 0.20, standardized mean difference (smd) = -0.18)), we found a large reduction in daytime waking activity in *arfgap2* mutants (**Fig. 2L-M**) (mean difference (95% CI) -0.90 s/awake minute (−1.21, -0.59), *P* <0.0001, smd = -1.34) indicative of an impaired movement phenotype.

**Fig. 2.**
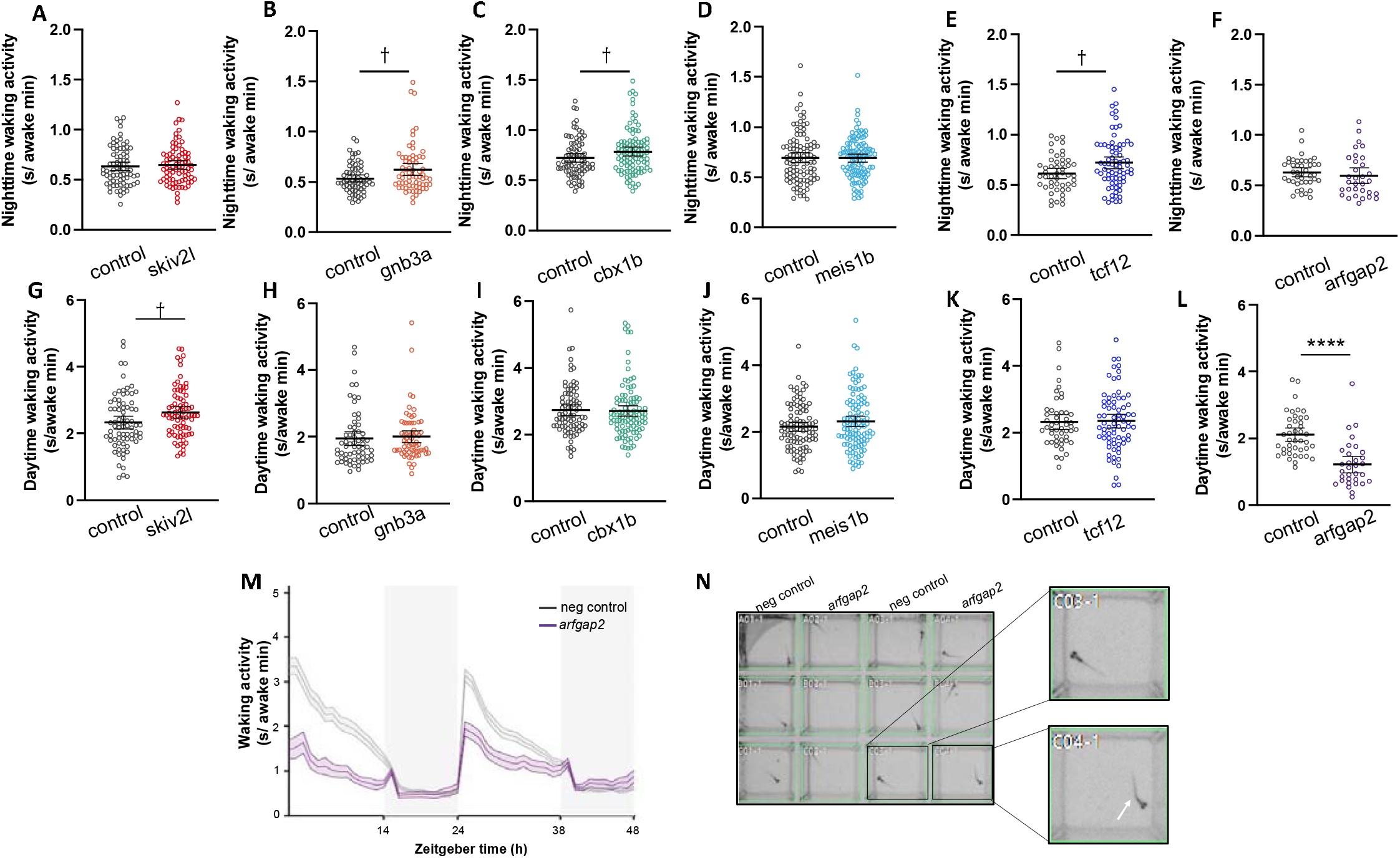
Waking activity and development is disrupted by mutation of *arfgap2*. **A-F**. Nighttime waking activity displayed as mean ± 95% CI for all mutants. **G-L**. Daytime waking activity displayed as mean ± 95% CI for all mutants. **M**. Waking activity graph for *arfgap2* mutants displayed as mean (line) and SEM (shaded). Gray shaded regions represent lights-off period. **N**. Representative image of *arfgap2* mutants and controls. Notable tail curvature (white arrow) is apparent in nearly all *arfgap2* mutants. ****p<0.0001 determined by Wilcoxon rank sum test and Hochberg step-up procedure for multiple comparisons (see **Methods**). † indicates significant (p < 0.05) before multiple comparisons. (n = 77 neg. control, 80 *skiv2l*); (n = 64 neg. control, 67 *gnb3a*); (n = 83 neg. control, 102 *cbx1b*); (n = 100 control, 111 *meis1b*); (n = 49 neg. control, 73 *tcf12*); (n = 42 control, 32 *arfgap2*).

Genes influencing insomnia are also commonly involved in neuronal development and can differentially influence behavior across developmental stages. Larval zebrafish develop an intact nervous system within the first few days of development and gene mutations that disrupt neurodevelopment lead to apparent changes in body morphology (Tran and Prober, 2022). We observed no gross morphological changes to *skiv2l, gnb3a, cbx1b, meis1b*, or *tcf12* mutants (data not shown); however, a clear and consistent morphological abnormality in *arfgap2* mutants was apparent beginning on approximately day 3 post fertilization, whereby nearly all mutants presented with a curvature in the tail (**Fig. 2N**). This morphological change paired with the markedly reduced waking activity suggests *arfgap2* is important during early development for proper motor development. Given this, we cannot reliably assess sleep/wake behavior in mutants with knockout of this gene.

### 2.2 Diurnal activity patterns are impacted by insomnia-associated genes

We next sought to describe the diurnal activity patterns in each of the mutants to determine whether insomnia-associated genes influence patterns of rest and activity. Insomnia complaints are often described as states of hypervigilance (Chen *et al*., 2014) and hyperarousal (Fernandez-Mendoza *et al*., 2016; Kalmbach *et al*., 2018). To capture similar states relating to hyperactivity in zebrafish, we measured activity patterns across light and dark periods. As expected, activity patterns showed robust entrainment by light-dark cycles in those mutants (**Fig. 3A-C**). Our previous work using neuron-specific RNAi resulted in significantly reduced activity duration in *Drosophila* (Palermo *et al*., 2021). In zebrafish, however, we observed significantly increased daytime activity duration in *skiv2l* mutants (mean difference (95% CI) = 20.56 s/h (5.61, 35.51), *P* = 0.005, smd = 0.43) (**Fig. 3D**) with no change in night activity mean difference (95% CI) = 1.97 s/h (−1.53, 5.46), P = 0.28, smd = 0.18) (**Fig. 3E**). *MEIS1* has commonly been associated with insomnia (Jansen *et al*., 2019; Watanabe *et al*., 2022) and restless leg syndrome (Lam *et al*., 2022; Salminen *et al*., 2017; Schulte *et al*., 2014; Spieler *et al*., 2014), and knockdown in *Drosophila* resulted in reduced sleep with no change to activity(Palermo *et al*., 2021). Likewise, in zebrafish, we observed no significant change to daytime activity (mean difference (95% CI) = 10.93 s/h (−1.62, 23.48), *P* = 0.2, smd = 0.23) (**Fig. 3F**) or nighttime activity (mean difference (95% CI) = 1.37 s/h (−2.21, 4.94), *P* = 0.25, smd = 0.10) (**Fig. 3G**). While knockdown of the *Drosophila* ortholog of *GNB3* displayed markedly reduced activity(Palermo *et al*., 2021), CRISPR mutation of the zebrafish ortholog did not present with altered activity (**Supplementary Fig. 1A and B)**, nor did *cbx1b* or *tcf12* mutants (**Supplementary Fig. 1C-F**).

**Fig. 3.**
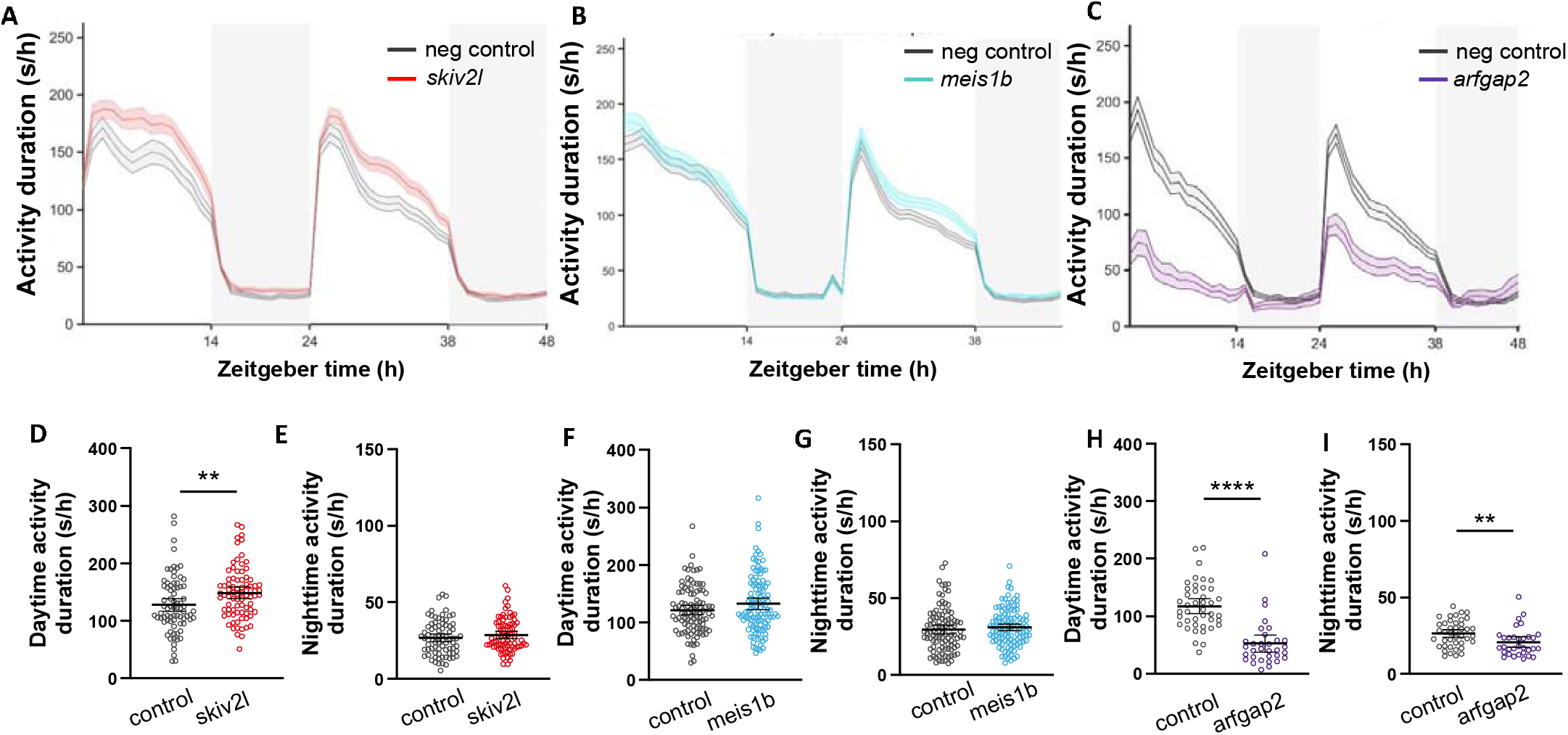
Activity duration is altered by *skiv2l* and *arfgap2* mutations. **A-C**. Rest-activity graphs for CRISPR mutants represented as mean (line) ± SEM (shaded) for activity duration (s/h). Shaded regions represent lights-off period. **D-E**. Mean ± 95% CI for day (**D**) and night (**E**) activity duration in *skiv2l* mutants (n = 77 neg control, 80 *skiv2l*). **F-G**. Mean ± 95% CI for day (F) and night (G) activity duration in *meis1b* mutants (n = 100 control, 111 meis1b). **H-I**. Mean ± 95% CI for day (**H**) and night (**I**) activity duration in arfgap2 mutants (n = 42 control, 32 arfgap2). Significance determined by Wilcoxon rank sum test followed by Hochberg step-up procedure for multiple comparisons (see **Methods**). *p<0.05, **p<0.01, ***p<0.001, ****p<0.0001.

Given the robust reduction in daytime waking activity observed in *arfgap2* mutants, we anticipated a reduction in total activity measured. Indeed, *arfgap2* mutants displayed significantly reduced daytime activity (mean difference (95% CI) = -64.92 (−83.98, -45.87), *P* < 0.0001, smd = -1.60) (**Fig. 3H**) and nighttime activity (mean difference (95% CI) = -5.62 (−9.81, - 1.43), *P* = 0.004, smd = -0.62) (**Fig. 3I**), demonstrating mutation of *arfgap2* greatly impairs movement in zebrafish.

### 2.3 Total sleep duration and latency to sleep onset is perturbed in CRISPR mutants

After screening for developmental and activity phenotypes, we characterized sleep in the six zebrafish mutants to determine if these insomnia-associated genes influenced sleep duration in vertebrates (zebrafish) in a similar manner to invertebrates (*Drosophila)*. We measured sleep across the fourteen-hour day and the ten-hour night using a standardized criterion of inactivity bouts lasting one minute or longer, as this has reliably been associated with the characteristics observed in mammalian sleep (e.g. elevated arousal threshold) (Chiu and Prober, 2013; Prober *et al*., 2006; Singh *et al*., 2015; Tran and Prober, 2022; Zhdanova *et al*., 2001). Diurnal sleep-wake patterns were intact in mutant and control fish (**Fig. 4A-B and Supplementary Fig. 3A**). Drosophila knockdown of the *SKIV2L* ortholog resulted in a robust increase in sleep duration (Palermo *et al*., 2021), but loss of *skiv2l* in zebrafish significantly reduced daytime sleep duration (mean difference (95% CI) = -5.11 minutes/hour (−7.77, -2.45), *P* = 0.0001, smd = - 0.61) (**Fig. 4C**), with a modest reduction in nighttime sleep duration (mean difference (95% CI) = -2.65 minutes/hour (−5.68, 0.37), *P* = 0.07, smd = -0.28) (**Fig. 4D**).

**Fig. 4.**
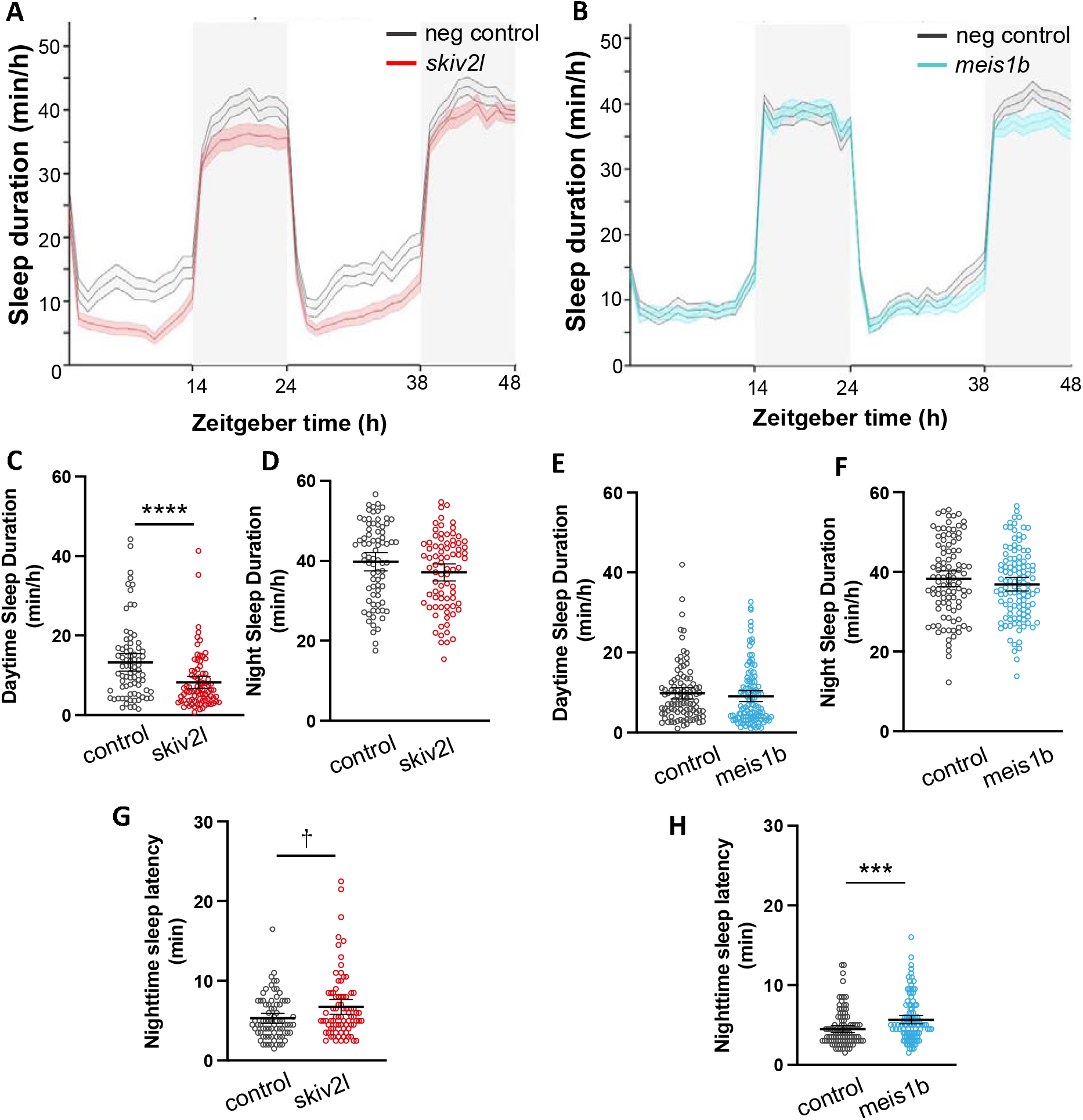
Total sleep duration in *skiv2l* and *meis1b* CRISPR mutants. Rest-activity graphs for *skiv2l* (**A**) and *meis1b* (**B**) mutants represented as mean (line) ± SEM (shaded) for sleep duration (min/h). Shaded regions represent lights-off period. Mean ± 95% CI for day (**C**) and night (**D**) sleep duration in *skiv2l* mutants (n = 77 neg. control, 80 *skiv2l*). Mean ± 95% CI for day (**E**) and night (**F**) sleep duration in *meis1b* mutants (n = 100 neg. control, 111 *meis1b*). Mean ± 95% CI for nighttime sleep latency in *skiv2l* (**G**) and *meis1b* (**H**) mutants. Significance determined by Wilcoxon rank sum test followed by Hochberg step-up procedure for multiple comparisons (see **Methods**). *p<0.05, **p<0.01, ***p<0.001, ****p<0.0001. † indicates significant (p < 0.05) before multiple comparisons.

Since waking activity and development were impacted by mutation of *arfgap2*, we cannot reliably assess sleep in these fish. Because our measurement of sleep is defined as bouts of inactivity greater than one minute, calculations of sleep appear to demonstrate an increase in both daytime (mean difference (95% CI) = 26.09 minutes/hour (20.52, 31.66), *P* < 0.0001, smd = 2.16) (**Supplementary Fig. 3B**) and nighttime sleep duration (mean difference (95% CI) = 8.37 minutes/hour (4.65, 12.09), *P* < 0.0001, smd = 1.06) (**Supplementary Fig. 3C**); however, this is likely an artifact caused by significantly reduced movement. These data demonstrate the importance of screening for developmental and activity phenotypes when relying on activity as a metric for sleep.

Despite finding a significant reduction in total sleep duration measured in *Drosophila* following knockdown of the *MEIS1 ortholog(Palermo et al*., *2021)*, loss of *meis1b* in zebrafish did not significantly (*P* > 0.05) alter total sleep duration (**Fig. 4E-F**). No significant changes in total sleep duration were observed in *gnb3a, tcf12*, and *cbx1b* mutants (**Supplementary Fig. 1G-L**).

Another key feature of insomnia is difficulty initiating sleep with an increase in latency to sleep onset. Since zebrafish sleep is tightly regulated by light-dark transition, a measure of sleep latency at night can be used to indicate time to sleep onset following lights-off. Mutations in *skiv2l* produced an increase in sleep latency (mean difference (95% CI) = 1.43 minutes (0.33, 2.52), *P* = 0.02, smd = 0.41) (**Fig. 4G**); however, this difference did not strictly meet the threshold for significance following multiple comparisons (see **Methods**). *Meis1b* mutants showed an increased sleep latency (mean difference (95%) = 1.13 minutes (0.44, 1.82), *P* = 0.0004, smd = 0.45) (**Fig. 4H**), supporting the role of this gene in insomnia-like behavior. Despite having reduced nighttime activity, *arfgap2* mutants did not have a significantly different sleep latency relative to controls (mean difference = -1.40 minutes (−2.56, -0.24), *P* = 0.08, smd = -0.54) (**Supplementary Fig. 3D**). Sleep latency was not altered in *gnb3a, tcf12*, or *cbx1b* mutants (**Supplementary Fig. 2A-C**).

### 2.4 Insomnia-associated genes contribute to changes in sleep continuity

Total sleep duration is not always impacted in patients with insomnia; rather, sleep is fragmented or considered not restorative leading to excessive daytime sleepiness (Association, 2013). We measured sleep bout length (the average length of each sleep period) during night and day as well as arousal threshold to observe changes to sleep depth across mutant lines. Furthermore, we measured sleep bout number (total number of sleep episodes) during day and night as a representation of sleep fragmentation. Consistent with reduced daytime sleep in *skiv2l* mutants, these larvae also demonstrated a reduction in sleep bout number during the day (mean difference (95% CI) = -1.75 bouts/hour (−2.61, -0.89), *P* = 0.0001, smd = -0.65) (**Fig. 5A**), with no change in bout number at night (mean difference (95% CI) = 0.26 bouts/hour (−0.39, 0.91), *P* = 0.52, smd = 0.13) (**Fig. 5B**). Although *meis1b* mutants did not present with a generalized sleep duration abnormality, they did demonstrate a significant increase in the number of sleep bouts at night (mean difference (95% CI) = 1.34 (0.80, 1.87) bouts/hour, *P* < 0.001, smd = 0.67) (**Fig. 5D**), with no change during the day (mean difference (95% CI) = 0.08 bouts/hour (−0.73, 0.90), *P* = 0.27, smd = 0.027) (**Fig. 5C**), indicating nighttime-specific sleep fragmentation caused by a gene that is commonly associated with RLS (El Gewely *et al*., 2018; Schulte *et al*., 2014).

**Fig. 5.**
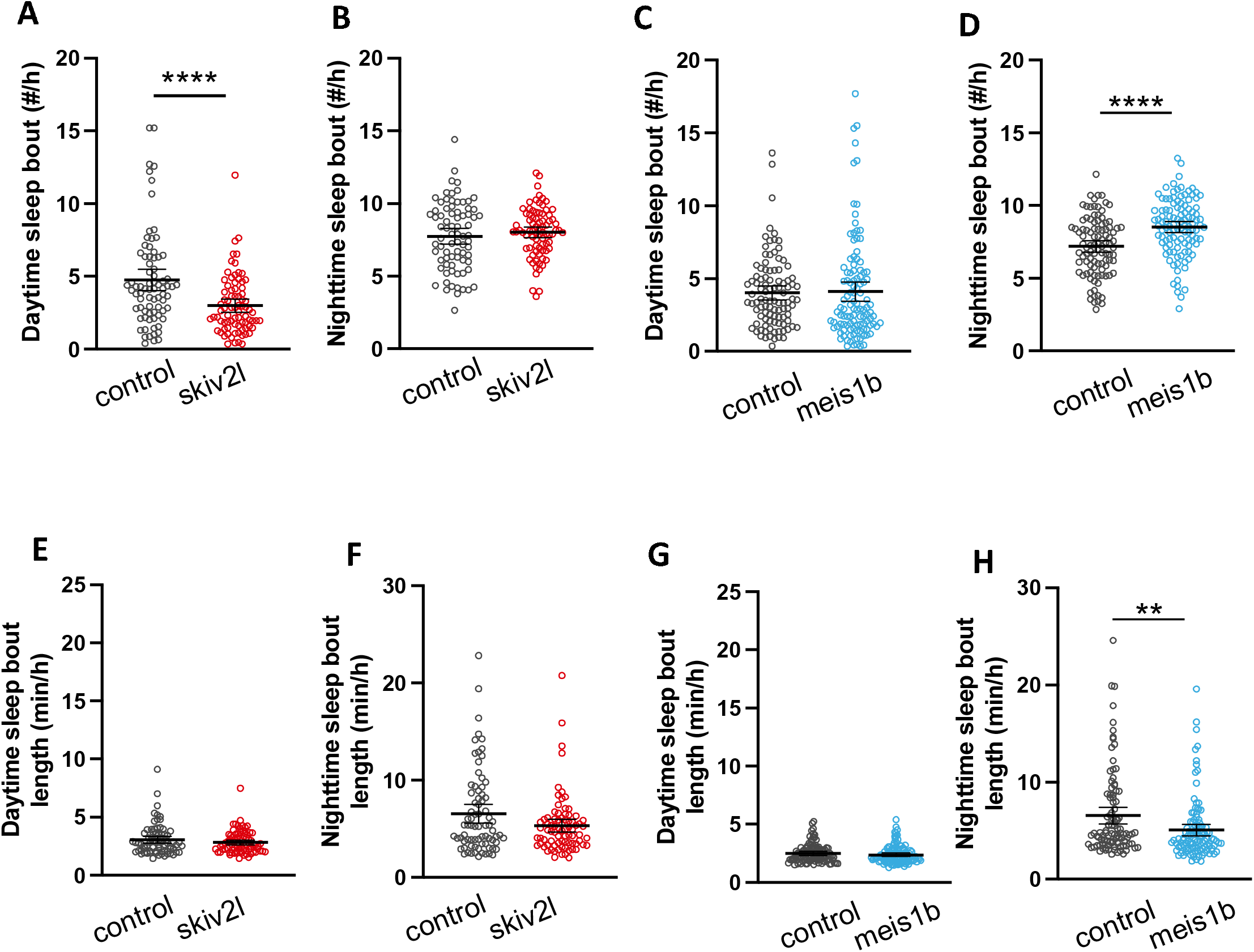
Sleep characteristics are significantly altered by CRISPR mutation of *skiv2l* and *meis1b*. Mean ± 95% CI for day and night sleep bout number in *skiv2l* (**A-B**) and *meis1b* (**C-D**) mutants. Mean ± 95% CI for day and night sleep bout length for *skiv2l* (**E-F**) and *meis1b* (**G-H**) mutants. Significance determined by Wilcoxon rank sum test followed by Hochberg step-up procedure for multiple comparisons (see **Methods**). *p<0.05, **p<0.01, ***p<0.001, ****p<0.0001. (n = 77 neg. control, 80 *skiv2l*); (n = 100 control, 111 *meis1b*).

There was no significant change in sleep bout length observed for *skiv2l* mutants during the day (mean difference (95% CI) = -0.22 minutes/bout (−0.58, 0.14), *P* = 0.51, smd = -0.20) or night (mean difference (95% CI) = -1.22 minutes/bout (−2.39, -0.05), P = 0.20, smd = -0.33 (**Fig. 5E and F**). Consistent with a fragmented sleep phenotype, nighttime sleep bout length was significantly shortened in *meis1b* mutants (mean difference (95% CI) = -1.49 minutes/bout (−2.52, -0.46), *P* = 0.002, smd = -0.4) (**Fig. 5H**), with no changes observed during the day (mean difference (95% CI) = -0.12 minutes/bout (−0.33, 0.10), *P* = 0.21, smd = -0.15) (**Fig. 5G**), demonstrating depth of sleep at night is also impacted by this gene.

We measured sleep bout number and bout length in *arfgap2* mutants to identify fragmented activity patterns throughout the day and night. Bouts of prolonged inactivity were apparent in *arfgap2* mutants manifesting as an increase in the number of daytime “sleep” bouts (mean difference (95% CI) = 4.11 bout/hour (2.39, 5.83), *P* < 0.0001, smd = 1.12) (**Supplementary Fig. 3E**). No change was observed for nighttime sleep bout number in *arfgap2* mutants (**Supplementary Fig. 3F**). Both day (mean difference = 2.85 minutes/bout (1.46, 4.23), *P* < 0.0001, smd = 1.04) (**Supplementary Fig. 3G**) and night inactivity bouts were longer (mean difference (95% CI) = 3.68 minutes/bout (1.25, 6.11), *P* = 0.002, smd = 0.75) (**Supplementary Fig. 3H**). These data imply that while *arfgap2* mutants spend the majority of their day in an immobile state, they do frequently switch between states of complete immobility and activity, as marked by an increase in the number of daytime “sleep” bouts.

Despite having no changes to total sleep or activity duration, *tcf12* mutants had fewer sleep bouts during the night (mean difference (95% CI) = -0.98 bouts/h (−1.74, -0.22), *P* = 0.005, smd = -0.47) (**Supplementary Fig. 4F**) and shorter sleep bouts during the night (mean difference (95.% CI) = -1.14 minutes/bout (−2.39, 0.10, *P* = 0.03, smd = -0.34)) (**Supplementary Fig. 4L**); however, these differences did not meet the strict threshold for significance following multiple comparisons (see **Methods**). No significant changes were observed for these sleep characteristics in the *gnb3a* or *cbx1b* mutants (**Supplementary Fig. 5A-D and G-J**). Together, these data support a conserved role for *meis1b* and *skiv2l* in promoting sleep-wake disruption primarily through altering sleep duration and consolidation. Although, the measured sleep characteristics in *tcf12* mutants did not meet the conservative threshold for significance following multiple comparisons, loss of *tcf12* does appear to impact nighttime sleep continuity.

Arousal threshold is increased during sleep in zebrafish (Prober *et al*., 2006; Zhdanova *et al*., 2001) similar to humans, and increased nighttime arousal is a common feature of insomnia (Bonnet and Arand, 2000; Mahowald and Schenck, 2005). Using mechano-acoustic stimuli of different intensities (**Supplementary Fig. 5B**), we measured the arousal response of each mutant line during the night. We measured EC50 for each mutant line and their respective control. EC50 compares the half-maximal response and corresponds to the stimulus intensity which elicits a response half-way between the minimum and maximum response and represents a threshold at which approximately half of the fish are aroused. While there is a slight shift to the right in the response curves at lower frequencies for *skiv2l* (**Supplementary Fig. 5C)**, *meis1b* **(Supplementary Fig. 5D)**, and *arfgap2* mutants **(Supplementary Fig. 5E)**, suggesting reduced arousal response at low stimuli intensities, the EC50 and maximal responses did not significantly differ for any group (*P* > 0.05 by extra sum-of-squares F-test). There was an expected step-wise increase in the fraction of responsive larvae as stimulus intensity increased (**Supplementary Fig. 5C-E**) indicating the majority of fish were asleep pre-stimulus and had an intact arousal response.

## 3. Discussion

There is an abundance of genomic data available through public repositories generated from GWAS and other sequencing approaches; however, functional characterization lags in validating the actual underlying genomic factors contributing to different phenotypes. Here, we demonstrate a proof-of-principle approach for moving from GWAS-implicated effector genes to validation in a vertebrate model organism for insomnia.

We elected to perform functional validation of top candidate genes identified using 3D genomics and *Drosophila* sleep data (Palermo *et al*., 2021). The candidate genes screened in this study have been mapped to insomnia GWAS-associated loci using ATAC-seq and promoter-focused Capture C protocols to identify high-resolution contacts between insomnia GWAS signals and effector genes (Palermo *et al*., 2021). Through a large-scale neuron-specific RNAi screen in adult *Drosophila melanogaster*, loss of function of these genes was shown to produce robust sleep phenotypes (Palermo *et al*., 2021), demonstrating their high conservation and potential regulatory function in sleep. The studies reported here tested the evolutionary conservation of function related to six genes which are highly conserved at the amino acid level across species and produced strong sleep phenotypes in *Drosophila* (*MEIS1, CBX1, TCF12, ARFGAP2, SKIV2L*, and *GNB3)*.

Increasingly, studies have shown that in addition to total sleep duration, altered sleep characteristics and poor sleep quality are predictive of negative health sequelae (Fernandez-Mendoza, 2017; Martin *et al*., 2011; Wallace *et al*., 2018). This includes day-to-day variability in sleep duration, sleep onset and waking time, as well as sleep fragmentation and excessive daytime sleepiness. Zebrafish provide a model system to observe these nuanced behaviors absent of external influences. Our results indicated disruptions to sleep continuity in multiple mutant lines as well as altered daytime sleep and activity, suggesting these genes play a role in more complex sleep-wake maintenance.

We observed that CRISPR mutation of *meis1b*, the strongest ortholog for human *MEIS1*, results in significantly fragmented nighttime sleep as well as increased sleep latency after lights-off. *MEIS1* is highly conserved, with 95% amino acid identity between zebrafish and humans(Hu *et al*., 2011). This gene encodes the Myeloid Ecotropic Viral Integration Site 1 protein, which acts as a transcription factor (Moskow *et al*., 1995). The *MEIS1* locus is one of the strongest association signals from previous insomnia GWAS (Jansen *et al*., 2019; Lane *et al*., 2017; Watanabe *et al*., 2022). Our variant 3D-mapping approach identified a putative causal variant in strong linkage disequilibrium (LD) with the sentinel GWAS SNP at this locus, rs1519102, which contacted the *MEIS1* promoter residing in open chromatin within NPCs (Palermo *et al*., 2021). rs1519102 resides within an intronic region, which suggests it is harbored in a cis-regulatory element acting as a transcriptional enhancer in a cell-specific manner(Lam *et al*., 2022). Work by Lam and colleagues (Lam *et al*., 2022) further identified expression quantitative trait loci (eQTL) residing within this region specific to brain cell types, including within the cerebellum, one of the brain regions where *MEIS1* is highly expressed. While *MEIS1* has repeatedly been identified as a candidate gene for insomnia (Hammerschlag *et al*., 2017; Jansen *et al*., 2019; Lane *et al*., 2017), there is debate as to whether its role in sleep disturbance is primarily due to its association with RLS (El Gewely *et al*., 2018; Watanabe *et al*., 2022). The consistent finding of *MEIS1* in GWAS for insomnia may represent a large proportion of undiagnosed RLS in the UK Biobank sample from which these data are derived (El Gewely *et al*., 2018). *MEIS1* knockouts have shown hyperactive phenotypes in mice (Salminen *et al*., 2017; Spieler *et al*., 2014), but it is unclear if this phenotype translates to sleep. Our model reveals a phenotype indicative of fragmented sleep that predominantly occurs at night, which is in line with the potential role in RLS, and implies that MEIS1 is acting in a circadian pattern to alter arousal and sleep consolidation. These mutants also had an increase in nighttime sleep latency, which is a common characteristic of insomnia.

Mutation of *skiv2l* in zebrafish significantly reduced sleep duration and had a modest effect in increasing sleep latency. In addition, these mutants showed daytime hyperactivity, suggesting a state of hyperarousal consistent with an insomnia-like condition. *SKIV2L* encodes the Superkiller Viralicilic Activity 2-Like RNA helicase and was 3D-mapped to the *MICB* GWAS locus (sentinel SNP rs3131638) on chromosome 6 (Palermo *et al*., 2021). It is located within a conserved region of the major histocompatibility complex (Dangel *et al*., 1995; Sultmann *et al*., 2000) and is believed to play a role in antiviral activity by blocking translation of poly(A) deficient mRNAs (Dangel *et al*., 1995). *SKIV2L* is fairly highly conserved between zebrafish and humans with 60% overall amino acid conservation and >90% conservation of the specific sequence encoding the DEAD box helicase (Hu *et al*., 2011), suggesting the role in RNA metabolism is particularly crucial. The mechanism by which *SKIV2L* acts to influence sleep is unknown; however, RNA metabolism is circadian-regulated and disturbed sleep has been shown to be associated with alterations in RNA metabolism (Moller-Levet *et al*., 2013). Loss-of-function in *SKIV2L* produced robust sleep phenotypes in both flies (Palermo *et al*., 2021) and zebrafish, suggesting strong conservation of function related to this gene. Interestingly, the sleep duration phenotype was opposite in these two model organisms. Although the *SKIV2L* locus is conserved, the GWAS variant (intron of MICB) is not. This suggests that while *SKIV2L* acts to regulate sleep in both species, its interaction with regulatory factors within each species may differentially modulate the behavior.

*ARFGAP2* encodes the ADP Ribosylation Factor GTPase Activating Protein 2 and was 3D-mapped to the *NDUFS3* GWAS locus (sentinel SNP rs11605348) on chromosome 11 (Palermo *et al*., 2021). It is 66% conserved at the amino acid level between human and zebrafish (Hu *et al*., 2011) and is highly expressed early in development (Alliance of Genome Resources, 2020), likely explaining the developmental phenotype observed in our larval model.

*ARFGAP2* has not been extensively studied in the context of behavioral characteristics; however, it has been associated with synaptic plasticity (Colameo *et al*., 2021; Zhang *et al*., 2012) and neurocognitive disorders including depression(Nagel *et al*., 2018) and Alzheimer’s disease (Gouveia *et al*., 2022), which are both associated with disrupted sleep. The zebrafish harboring mutations in *arfgap2* appeared normal through early development (1-2 days post fertilization); however, by the third day, approximately half of the larvae began to appear abnormal with a curved tail. By 5 days post fertilization, when the sleep assay began, nearly all mutant larvae appeared developmentally abnormal; however, very few died. *Arfgap2* expression is high during this time frame and likely serves an important role during a critical period of development. Single cell RNA sequencing data show expression of *ARFGAP2* in skeletal myocytes (Karlsson *et al*., 2021), which may contribute to the morphological and movement abnormalities.

ATAC-seq and promoter-focused capture C showed proxy SNPs at the *TCF12* insomnia GWAS locus contacting its own promoter (Palermo *et al*., 2021). Human data indicate *TCF12* is highly expressed in oligodendrocytes and their precursors (Karlsson *et al*., 2021) and controls oligodendroglial cell proliferation through transcriptional regulation (Wang *et al*., 2014), which has been implicated in sleep regulation (Bellesi, 2015; Bellesi *et al*., 2013). Although the sleep traits we measured in *tcf12* mutants showed modest effects (smd 0.34-0.48) that did not meet the significance threshold following multiple comparisons, their sleep was consistently abnormal across multiple nighttime parameters including nighttime waking activity, sleep bout number, and sleep bout length at night, indicating this transcription factor may play a role in night-specific activity regulation similar to *meis1*.

Surprisingly, we did not observe a significant phenotype in *gnb3a* mutants. This gene has been shown to be associated with sleep quality and diurnal preference in humans (Parsons *et al*., 2014) and is highly conserved in vertebrates (Ritchey *et al*., 2010). Single cell RNA sequencing data indicate *GNB3* is most highly expressed in retinal cells (Karlsson *et al*., 2021) and is associated with congenital stationary night blindness (Vincent *et al*., 2016). While *gnb3* may be important for sensing changes to light that may influence diurnal activity regulation, zebrafish larvae have other photoreceptive cells that may compensate for loss of this gene (Fernandes *et al*., 2012).

We did not observe significant changes to sleep or activity in *cbx1* mutant zebrafish. Our 3D genomics data identified multiple genes at this GWAS locus on chromosome 17 contacted by the insomnia-associated SNPs (Palermo *et al*., 2021); therefore, this gene may not act independently in vertebrates to influence sleep-wake behaviors.

## 4. Limitations

The current work was based on GWAS data primarily from individuals of European ancestry and may not be generalizable across all ancestral groups.

Several of these genes exhibit differential expression across development. In these experiments, we assayed larval zebrafish, while, in contrast, our previous work tested sleep in adult *Drosophila*. Knockdown of the *arfgap2* ortholog in *Drosophila* produced a significant reduction in sleep duration, yet the CRISPR mutation in zebrafish produced abnormal wake and sleep patterns paired with developmental abnormalities. The differences in phenotypes observed between larval zebrafish and adult *Drosophila* with perturbed *arfgap2* expression is likely due to developmental expression differences, demonstrating the benefit of cross-species and cross-development observation. Moreover, our experiments in *Drosophila* (Palermo *et al*., 2021) used neuron-specific RNA-interference to knockdown gene expression in neuronal cells, sparing expression elsewhere. This, too, likely contributed to the phenotypic differences observed.

Using F0 larvae for screening is a rapid and efficient approach for assaying sleep behavior to narrow long lists of candidate genes (Kroll *et al*., 2021); however, it has inherent limitations. While we found that extremely little mosaicism was apparent in our larvae harboring the CRISPR mutations, there was a high degree of variability in behavior of both mutants and controls. The variability is possibly due to trauma induced by the injections, which is why we used scramble-sgRNA-injected controls for comparison. This variability likely diminishes true sleep-activity phenotypes and may result in false-negative results. Given we previously observed robust sleep phenotypes in *Drosophila* RNAi lines for these genes (Palermo *et al*., 2021), we cannot rule out those candidate genes that demonstrated minimal phenotypes; however this approach highlights those with particularly strong phenotypes in a vertebrate model. Rather, future studies assessing stable F1 mutant lines represent a promising tool to assess these genes. Additionally, several of these genes are duplicated in zebrafish and one ortholog may offer compensatory action over the mutated gene. We chose to focus on the ortholog with strongest conservation across species as we hypothesized that this would be most consistent with the *Drosophila* screen; however, double knockouts are warranted in the future to assess behavior.

## 5. Conclusion

The genes examined in the current study were implicated as putative causal genes associated with insomnia GWAS signals through 3D genomics approaches and were shown to significantly impact invertebrate sleep characteristics indicating high conservation of function. We demonstrate that the orthologs of *SKIV2L* and *MEIS1* are deeply conserved and important for vertebrate sleep maintenance as well. We also find that *ARFGAP2* is required for proper development of motor behaviors to produce normal activity rhythms. This work also demonstrates the utility of employing cross-species paradigms to examine conserved behaviors, as we show that not all genes have strong conservation of function across different model organisms. Together, we provide rationale for the functional interrogation of GWAS-associated effector genes using large-scale screening approaches in model organisms to identify promising target genes for disease intervention.

## 6. Methods

### 6.1 Animal use

All experiments with zebrafish were conducted in accordance with the University of Pennsylvania Institutional Animal Care and Use Committee guidelines. Breeding pairs consisted of wild-type AB and TL (Tupfel Long-fin) strains. Fish were housed in standard conditions with 14-hour:10-hour light:dark cycle at 28.5°C, with lights on at 9 a.m (ZT0). and lights off at 11 p.m (ZT14).

### 6.2 CRISPR/Cas9 mutagenesis

Single guide RNAs (sgRNAs) were designed using the online tool Crispor (http://crispor.tefor.net/) with the reference genome set to “NCBI GRCz11” and the protospacer adjacent motif (PAM) set to “20bp-NGG-Sp Cas9, SpCas9-HF, eSpCas9 1.1.” sgRNAs were prioritized by specificity score (>95%) with 0 predicted off-targets with up to 3 mismatches. The zebrafish sequence was obtained using Ensembl (https://useast.ensembl.org/) with GRCz11 as the reference genome. Sequence was aligned to the human amino acid sequence using MARRVEL (http://marrvel.org/) to identify the region with highest conservation, and each sgRNA was designed targeting this conserved exonic region (**Supplementary Table 2**). AB/TL breeding pairs were set up overnight and embryos collected in embryonic growth media (E3 medium; 5mM NaCl, 0.17 mM KCl, 0.33 mM CaCl2, 0.33 mM MgSO4) the following morning shortly after lights-on. Pre-formed ribonuclear protein (RNP) complexes containing the sgRNA and Cas9 enzyme were injected at the single-cell stage alternating between the gene group and scramble-injected negative control group. Embryos were left unperturbed for one day before being transported to fresh E3 media in petri dishes (approximately 50 per dish). All embryos and larvae were housed in an incubator at 28.5°C, with lights on at 9 a.m. (ZT0) and lights off at 11 p.m (ZT14). Dead embryos and chorion membranes were removed daily until day 5 post fertilization. On day 5, CRISPR mutants and scramble-injected controls were pipetted into individual wells of a 96-well plate and placed into a zebrabox (Viewpoint Life Sciences) for automated video monitoring. Genotypes were placed into alternating rows to minimize location bias within the plate. Each zebrabox is sound-attenuating and contains circulating water held at a temperature of 28.5ºC with automated lights cycling on the same 14-hour:10-hour light/dark cycle. Sleep-wake behaviors were measured through automated video-tracking, as described previously (Kroll *et al*., 2021; Palermo *et al*., 2021; Rihel *et al*., 2010).

### 6.3 DNA extraction and PCR for genotyping

DNA extraction was performed per the manufacturer’s protocol (Quanta bio, Beverly, MA) immediately following completion of the sleep assay, as described previously(Palermo *et al*., 2021). Larvae were euthanized by rapid cooling on a mixture of ice and water between 2-4°C for a minimum of 30 minutes after complete cessation of movement was observed. Genotyping was performed on individual fish at the conclusion of each sleep assay. Either restriction digest or headloop PCR methods were used to validate mutations (Kroll *et al*., 2021; Palermo *et al*., 2021; Rand *et al*., 2005). Primers for genotyping are listed in **Supplementary Table 3**. All primers were run on a 2% agarose gel and sequence verified using Sanger sequencing to verify the target region.

### 6.4 Data collection and analysis for sleep phenotyping

Activity data were captured using automated video tracking (Viewpoint Life Sciences) software in quantization mode (Palermo *et al*., 2021). As described previously (Chen *et al*., 2017), threshold for detection was set as the following: detection threshold: 20; burst: 29; freeze: 3; bin size: 60 seconds. Data were processed using custom MATLAB scripts (Lee *et al*., 2022) to calculate the following parameters for both day and night separately: sleep duration (minutes/hour), activity duration (seconds/hour), waking activity (seconds/awake minute/hour), sleep bout length (minutes/bout), sleep bout number (number/hour) and nighttime sleep latency (minutes). All animals were allowed to acclimate to the zebrabox for approximately 24 hours before beginning continuous data collection for 48 hours starting at lights-on.

### 6.5 Arousal Threshold Assay

A mechano-acoustic stimulus was used to determine arousal threshold using a protocol adapted from previous work (Reichert *et al*., 2019; Singh *et al*., 2015). Individual fish were placed into alternating columns of a 96-well plate (**Supplementary Fig. 5A**) to avoid location bias. Ten different vibration frequencies were applied, which consistently produced a step-wise increase in arousability. Frequencies were pseudo-randomly assigned to prevent acclimation to any given stimulus frequency throughout the trials. Frequency steps of 40Hz were ordered as follows: 560Hz, 400Hz, 520Hz, 720Hz, 440Hz, 680Hz, 480Hz, 760Hz, 600Hz, 640Hz. These ten frequencies were each presented ten times for 5 seconds every 3 minutes (5 seconds on, 2 mins 55 seconds off) beginning at 1 a.m. (ZT16) and ending at 6 a.m. (ZT21) (**Supplementary Fig. 5B**). Lower frequencies produced larger changes in movement and an increase in the fraction of responsive larvae. Therefore, analyses were presented as highest-to-lowest frequency representing lowest-to-highest intensity of stimulation (i.e. 760Hz, 720Hz, 680Hz, 640Hz, 600Hz, 560Hz, 520Hz, 480Hz, 440Hz, 400Hz). The response to stimuli was measured by automated video tracking and analyzed using Matlab (Mathworks) and Excel (Microsoft). The response fraction represents the percentage of larvae considered to be asleep during the 5 second pre-stimulus baseline (activity <0.01s/5sec, determined empirically using average movement prestimulus) and whose activity increased over baseline during the stimulus presentation. Curve-fit and statistical analyses were carried out in Prism V9 (Graphpad) using non-linear regression (Agonist vs Response--Variable slope (four parameters)) with extra sum-of-squares F-test used to compare EC50 (half-maximal response) and top (maximal response) as described previously (Singh *et al*., 2015).

### 6.6 Statistical analysis and control for multiple comparisons across sleep traits

Continuous data are summarized using means and 95% confidence intervals (CI). Effect sizes are described as standardized mean difference (smd). Phenotypes of interest included a total of eleven measurements related to sleep and activity, including day and night sleep duration, activity, waking activity, sleep bout length, and sleep bout number, as well as nocturnal sleep latency. Analyses compared these traits between scramble-injected and CRIPSR mutant fish for six different candidate genes – *skiv2l, meis1b, gnb3a, cbx1b, tcf12*, and *arfgap2*. Primary comparisons between mutant and scramble-injected fish were performed using non-parametric Wilcoxon rank-sum tests to mitigate any potential impact of non-normality of the endpoints. A Hochberg step-up procedure (Hochberg, 1988; Huang and Hsu, 2007) was applied to the analysis of each gene to maintain gene-specific type I error at the desired level of 0.05 across the tested hypotheses. Briefly, to implement the Hochberg method the p-values for the set of eleven null hypotheses are ordered from largest to smallest, and each p-value is compared to a sequentially decreasing alpha-level to determine whether the associated null hypothesis (and subsequent hypotheses) should be rejected. Symbolically, for the set of p-values {p_1_, … p_11_}, ordered from largest to smallest and testing the corresponding set of null hypotheses {H_o1_, … H_o11_}, the procedure is implemented as:

**Step 1:** Evaluate whether p_1_ < 0.05. If yes, reject H_o1_ and all subsequent null hypotheses {H_o2_, … H_o11_}. Else, do not reject H_o1_ and go to Step 2.

**Step 2:** Evaluate whether p_2_ < 0.05/2. If yes, reject H_o2_ and all subsequent null hypotheses {H_o3_, … H_o11_}. Else, do not reject H_o2_ and go to Step 3. […]

**Step 11:** Evaluate whether p_k_ < 0.05/11. If yes, reject H_o11_. Else, none of the null hypotheses {H_o1_, … H_o11_} are rejected and stop.

Comparisons may be significant by the rank sum analysis but not reach the threshold for significance following multiple comparisons using the Hochberg approach described. In this case, results should be interpreted with caution.

## Acknowledgements

The work was supported by NIH grant T32 HL07953, R01 HL143790, and P01 HL094307. Dr. Grant is supported by NIH awards R01 AG057516 and R01 HD056465 and the Daniel B. Burke Endowed Chair.

## Author contributions

A.J.Z. designed and carried out experiments, analyzed data, and wrote and edited the manuscript. F.D.B. assisted with experiments and edited the manuscript. B.T.K. performed statistical analyses and edited the manuscript. Z.Y.S. assisted with experiments. J.P., A.C., S.S., M.C.P., E.B.B., J.A.P., A.D.W., O.J.V., D.R.M., and A.K., assisted with experimental design, data collection, interpretation, and edited the manuscript. P.R.G., A.C.K., S.F.G., and A.I.P conceptualized and designed experiments and contributed to writing and editing the manuscript.

## Graphical Abstract

Created using BioRender.

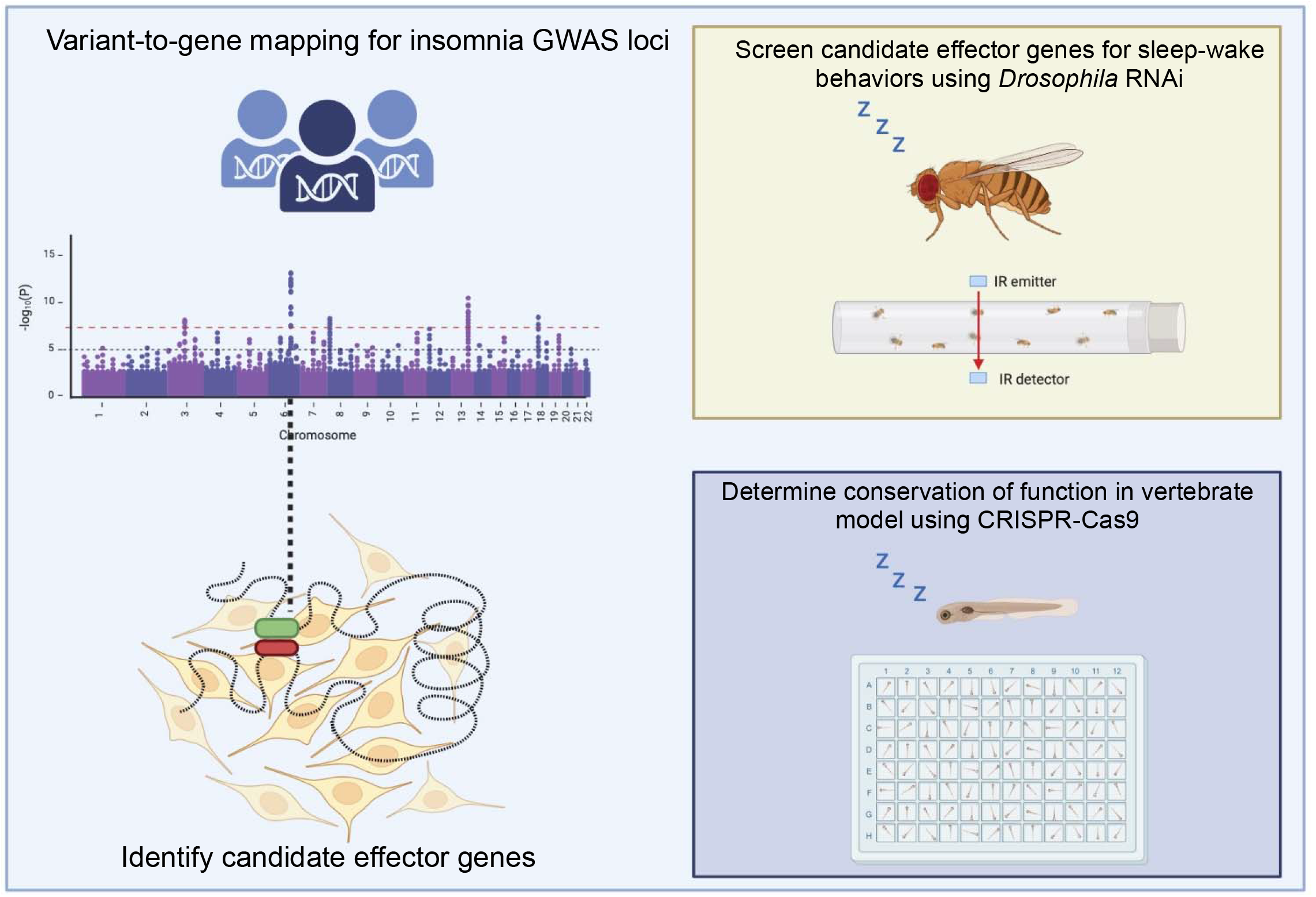

## References

Alliance of Genome Resources, C., 2020. Alliance of Genome Resources Portal: unified model organism research platform. Nucleic Acids Res 48, D650–D658.

Association, A.P., 2013. Diagnostic and statistical manual of mental disorders

Barlow, I.L., Rihel, J., 2017. Zebrafish sleep: from geneZZZ to neuronZZZ. Curr Opin Neurobiol 44, 65–71.

Bellesi, M., 2015. Sleep and oligodendrocyte functions. Curr Sleep Med Rep 1, 20–26.

Bellesi, M., Pfister-Genskow, M., Maret, S., Keles, S., Tononi, G., Cirelli, C., 2013. Effects of sleep and wake on oligodendrocytes and their precursors. J Neurosci 33, 14288–14300.

Bonnet, M.H., Arand, D.L., 2000. Activity, arousal, and the MSLT in patients with insomnia. Sleep 23, 205–212.

Chen, M.C., Chang, C., Glover, G.H., Gotlib, I.H., 2014. Increased insula coactivation with salience networks in insomnia. Biol Psychol 97, 1–8.

Chen, S., Reichert, S., Singh, C., Oikonomou, G., Rihel, J., Prober, D.A., 2017. Light-Dependent Regulation of Sleep and Wake States by Prokineticin 2 in Zebrafish. Neuron 95, 153–168 e156.

Chesi, A., Wagley, Y., Johnson, M.E., Manduchi, E., Su, C., Lu, S., Leonard, M.E., Hodge, K.M., Pippin, J.A., Hankenson, K.D., Wells, A.D., Grant, S.F.A., 2019. Genome-scale Capture C promoter interactions implicate effector genes at GWAS loci for bone mineral density. Nat Commun 10, 1260.

Chiu, C.N., Prober, D.A., 2013. Regulation of zebrafish sleep and arousal states: current and prospective approaches. Front Neural Circuits 7, 58.

Claussnitzer, M., Cho, J.H., Collins, R., Cox, N.J., Dermitzakis, E.T., Hurles, M.E., Kathiresan, S., Kenny, E.E., Lindgren, C.M., MacArthur, D.G., North, K.N., Plon, S.E., Rehm, H.L., Risch, N., Rotimi, C.N., Shendure, J., Soranzo, N., McCarthy, M.I., 2020. A brief history of human disease genetics. Nature 577, 179–189.

Colameo, D., Rajman, M., Soutschek, M., Bicker, S., von Ziegler, L., Bohacek, J., Winterer, J., Germain, P.L., Dieterich, C., Schratt, G., 2021. Pervasive compartment-specific regulation of gene expression during homeostatic synaptic scaling. EMBO Rep 22, e52094.

Dangel, A.W., Shen, L., Mendoza, A.R., Wu, L.C., Yu, C.Y., 1995. Human helicase gene SKI2W in the HLA class III region exhibits striking structural similarities to the yeast antiviral gene SKI2 and to the human gene KIAA0052: emergence of a new gene family. Nucleic Acids Res 23, 2120–2126.

Dashti, H.S., Jones, S.E., Wood, A.R., Lane, J.M., van Hees, V.T., Wang, H., Rhodes, J.A., Song, Y., Patel, K., Anderson, S.G., Beaumont, R.N., Bechtold, D.A., Bowden, J., Cade, B.E., Garaulet, M., Kyle, S.D., Little, M.A., Loudon, A.S., Luik, A.I., Scheer, F., Spiegelhalder, K., Tyrrell, J., Gottlieb, D.J., Tiemeier, H., Ray, D.W., Purcell, S.M., Frayling, T.M., Redline, S., Lawlor, D.A., Rutter, M.K., Weedon, M.N., Saxena, R., 2019. Genome-wide association study identifies genetic loci for self-reported habitual sleep duration supported by accelerometer-derived estimates. Nat Commun 10, 1100.

El Gewely, M., Welman, M., Xiong, L., Yin, S., Catoire, H., Rouleau, G., Montplaisir, J.Y., Desautels, A., Warby, S.C., 2018. Reassessing GWAS findings for the shared genetic basis of insomnia and restless legs syndrome. Sleep 41.

Fernandes, A.M., Fero, K., Arrenberg, A.B., Bergeron, S.A., Driever, W., Burgess, H.A., 2012. Deep brain photoreceptors control light-seeking behavior in zebrafish larvae. Curr Biol 22, 2042–2047.

Fernandez-Mendoza, J., 2017. The insomnia with short sleep duration phenotype: an update on it’s importance for health and prevention. Curr Opin Psychiatry 30, 56–63.

Fernandez-Mendoza, J., Li, Y., Vgontzas, A.N., Fang, J., Gaines, J., Calhoun, S.L., Liao, D., Bixler, E.O., 2016. Insomnia is Associated with Cortical Hyperarousal as Early as Adolescence. Sleep 39, 1029–1036.

Forgetta, V., Jiang, L., Vulpescu, N.A., Hogan, M.S., Chen, S., Morris, J.A., Grinek, S., Benner, C., Jang, D.K., Hoang, Q., Burtt, N., Flannick, J.A., McCarthy, M.I., Fauman, E., Greenwood, C.M.T., Maurano, M.T., Richards, J.B., 2022. An effector index to predict target genes at GWAS loci. Hum Genet 141, 1431–1447.

Freeman, A.A., Syed, S., Sanyal, S., 2013. Modeling the genetic basis for human sleep disorders in Drosophila. Commun Integr Biol 6, e22733.

Fulco, C.P., Nasser, J., Jones, T.R., Munson, G., Bergman, D.T., Subramanian, V., Grossman, S.R., Anyoha, R., Doughty, B.R., Patwardhan, T.A., Nguyen, T.H., Kane, M., Perez, E.M., Durand, N.C., Lareau, C.A., Stamenova, E.K., Aiden, E.L., Lander, E.S., Engreitz, J.M., 2019. Activity-by-contact model of enhancer-promoter regulation from thousands of CRISPR perturbations. Nat Genet 51, 1664–1669.

Gouveia, C., Gibbons, E., Dehghani, N., Eapen, J., Guerreiro, R., Bras, J., 2022. Genome-wide association of polygenic risk extremes for Alzheimer’s disease in the UK Biobank. Sci Rep 12, 8404.

Hammerschlag, A.R., Stringer, S., de Leeuw, C.A., Sniekers, S., Taskesen, E., Watanabe, K., Blanken, T.F., Dekker, K., Te Lindert, B.H.W., Wassing, R., Jonsdottir, I., Thorleifsson, G., Stefansson, H., Gislason, T., Berger, K., Schormair, B., Wellmann, J., Winkelmann, J., Stefansson, K., Oexle, K., Van Someren, E.J.W., Posthuma, D., 2017. Genome-wide association analysis of insomnia complaints identifies risk genes and genetic overlap with psychiatric and metabolic traits. Nat Genet 49, 1584–1592.

Hendricks, J.C., Sehgal, A., Pack, A.I., 2000. The need for a simple animal model to understand sleep. Prog Neurobiol 61, 339–351.

Hochberg, Y., 1988. A sharper bonferroni procedure for multiple tests of significance. New York University Graduate School of Business Administration.

Howe, K., Clark, M.D., Torroja, C.F., Torrance, J., Berthelot, C., Muffato, M., Collins, J.E., Humphray, S., McLaren, K., Matthews, L., McLaren, S., Sealy, I., Caccamo, M., Churcher, C., Scott, C., Barrett, J.C., Koch, R., Rauch, G.J., White, S., Chow, W., Kilian, B., Quintais, L.T., Guerra-Assuncao, J.A., Zhou, Y., Gu, Y., Yen, J., Vogel, J.H., Eyre, T., Redmond, S., Banerjee, R., Chi, J., Fu, B., Langley, E., Maguire, S.F., Laird, G.K., Lloyd, D., Kenyon, E., Donaldson, S., Sehra, H., Almeida-King, J., Loveland, J., Trevanion, S., Jones, M., Quail, M., Willey, D., Hunt, A., Burton, J., Sims, S., McLay, K., Plumb, B., Davis, J., Clee, C., Oliver, K., Clark, R., Riddle, C., Elliot, D., Threadgold, G., Harden, G., Ware, D., Begum, S., Mortimore, B., Kerry, G., Heath, P., Phillimore, B., Tracey, A., Corby, N., Dunn, M., Johnson, C., Wood, J., Clark, S., Pelan, S., Griffiths, G., Smith, M., Glithero, R., Howden, P., Barker, N., Lloyd, C., Stevens, C., Harley, J., Holt, K., Panagiotidis, G., Lovell, J., Beasley, H., Henderson, C., Gordon, D., Auger, K., Wright, D., Collins, J., Raisen, C., Dyer, L., Leung, K., Robertson, L., Ambridge, K., Leongamornlert, D., McGuire, S., Gilderthorp, R., Griffiths, C., Manthravadi, D., Nichol, S., Barker, G., Whitehead, S., Kay, M., Brown, J., Murnane, C., Gray, E., Humphries, M., Sycamore, N., Barker, D., Saunders, D., Wallis, J., Babbage, A., Hammond, S., Mashreghi-Mohammadi, M., Barr, L., Martin, S., Wray, P., Ellington, A., Matthews, N., Ellwood, M., Woodmansey, R., Clark, G., Cooper, J., Tromans, A., Grafham, D., Skuce, C., Pandian, R., Andrews, R., Harrison, E., Kimberley, A., Garnett, J., Fosker, N., Hall, R., Garner, P., Kelly, D., Bird, C., Palmer, S., Gehring, I., Berger, A., Dooley, C.M., Ersan-Urun, Z., Eser, C., Geiger, H., Geisler, M., Karotki, L., Kirn, A., Konantz, J., Konantz, M., Oberlander, M., Rudolph-Geiger, S., Teucke, M., Lanz, C., Raddatz, G., Osoegawa, K., Zhu, B., Rapp, A., Widaa, S., Langford, C., Yang, F., Schuster, S.C., Carter, N.P., Harrow, J., Ning, Z., Herrero, J., Searle, S.M., Enright, A., Geisler, R., Plasterk, R.H., Lee, C., Westerfield, M., de Jong, P.J., Zon, L.I., Postlethwait, J.H., Nusslein-Volhard, C., Hubbard, T.J., Roest Crollius, H., Rogers, J., Stemple, D.L., 2013. The zebrafish reference genome sequence and its relationship to the human genome. Nature 496, 498–503.

Hu, Y., Flockhart, I., Vinayagam, A., Bergwitz, C., Berger, B., Perrimon, N., Mohr, S.E., 2011. An integrative approach to ortholog prediction for disease-focused and other functional studies. BMC Bioinformatics 12, 357.

Huang, Y., Hsu, J.C., 2007. Hochberg’s Step-up Method: Cutting Corners off Holm’s Step-down Method. Biometrika 94, 965–975.

Jansen, P.R., Watanabe, K., Stringer, S., Skene, N., Bryois, J., Hammerschlag, A.R., de Leeuw, C.A., Benjamins, J.S., Munoz-Manchado, A.B., Nagel, M., Savage, J.E., Tiemeier, H., White, T., andMe Research, T., Tung, J.Y., Hinds, D.A., Vacic, V., Wang, X., Sullivan, P.F., van der Sluis, S., Polderman, T.J.C., Smit, A.B., Hjerling-Leffler, J., Van Someren, E.J.W., Posthuma, D., 2019. Genome-wide analysis of insomnia in 1,331,010 individuals identifies new risk loci and functional pathways. Nat Genet 51, 394–403.

Kalmbach, D.A., Cuamatzi-Castelan, A.S., Tonnu, C.V., Tran, K.M., Anderson, J.R., Roth, T., Drake, C.L., 2018. Hyperarousal and sleep reactivity in insomnia: current insights. Nat Sci Sleep 10, 193–201.

Karlsson, M., Zhang, C., Mear, L., Zhong, W., Digre, A., Katona, B., Sjostedt, E., Butler, L., Odeberg, J., Dusart, P., Edfors, F., Oksvold, P., von Feilitzen, K., Zwahlen, M., Arif, M., Altay, O., Li, X., Ozcan, M., Mardinoglu, A., Fagerberg, L., Mulder, J., Luo, Y., Ponten, F., Uhlen, M., Lindskog, C., 2021. A single-cell type transcriptomics map of human tissues. Sci Adv 7.

Kroll, F., Powell, G.T., Ghosh, M., Gestri, G., Antinucci, P., Hearn, T.J., Tunbak, H., Lim, S., Dennis, H.W., Fernandez, J.M., Whitmore, D., Dreosti, E., Wilson, S.W., Hoffman, E.J., Rihel, J., 2021. A simple and effective F0 knockout method for rapid screening of behaviour and other complex phenotypes. Elife 10.

Lam, D.D., Antic Nikolic, A., Zhao, C., Mirza-Schreiber, N., Krezel, W., Oexle, K., Winkelmann, J., 2022. Intronic elements associated with insomnia and restless legs syndrome exhibit cell-type-specific epigenetic features contributing to MEIS1 regulation. Hum Mol Genet 31, 1733–1746.

Lane, J.M., Liang, J., Vlasac, I., Anderson, S.G., Bechtold, D.A., Bowden, J., Emsley, R., Gill, S., Little, M.A., Luik, A.I., Loudon, A., Scheer, F.A., Purcell, S.M., Kyle, S.D., Lawlor, D.A., Zhu, X., Redline, S., Ray, D.W., Rutter, M.K., Saxena, R., 2017. Genome-wide association analyses of sleep disturbance traits identify new loci and highlight shared genetics with neuropsychiatric and metabolic traits. Nat Genet 49, 274–281.

Lappalainen, T., MacArthur, D.G., 2021. From variant to function in human disease genetics. Science 373, 1464–1468.

Lasconi, C., Pahl, M.C., Cousminer, D.L., Doege, C.A., Chesi, A., Hodge, K.M., Leonard, M.E., Lu, S., Johnson, M.E., Su, C., Hammond, R.K., Pippin, J.A., Terry, N.A., Ghanem, L.R., Leibel, R.L., Wells, A.D., Grant, S.F.A., 2021. Variant-to-Gene-Mapping Analyses Reveal a Role for the Hypothalamus in Genetic Susceptibility to Inflammatory Bowel Disease. Cell Mol Gastroenterol Hepatol 11, 667–682.

Lasconi, C., Pahl, M.C., Pippin, J.A., Su, C., Johnson, M.E., Chesi, A., Boehm, K., Manduchi, E., Ou, K., Golson, M.L., Wells, A.D., Kaestner, K.H., Grant, S.F.A., 2022. Variant-to-gene-mapping analyses reveal a role for pancreatic islet cells in conferring genetic susceptibility to sleep-related traits. Sleep 45.

Lee, D.A., Oikonomou, G., Prober, D.A., 2022. Large-scale Analysis of Sleep in Zebrafish. Bio Protoc 12, e4313.

Mahowald, M.W., Schenck, C.H., 2005. Insights from studying human sleep disorders. Nature 437, 1279–1285.

Martin, J.L., Fiorentino, L., Jouldjian, S., Mitchell, M., Josephson, K.R., Alessi, C.A., 2011. Poor self-reported sleep quality predicts mortality within one year of inpatient post-acute rehabilitation among older adults. Sleep 34, 1715–1721.

Moller-Levet, C.S., Archer, S.N., Bucca, G., Laing, E.E., Slak, A., Kabiljo, R., Lo, J.C., Santhi, N., von Schantz, M., Smith, C.P., Dijk, D.J., 2013. Effects of insufficient sleep on circadian rhythmicity and expression amplitude of the human blood transcriptome. Proc Natl Acad Sci U S A 110, E1132–1141.

Morin, C.M., Drake, C.L., Harvey, A.G., Krystal, A.D., Manber, R., Riemann, D., Spiegelhalder, K., 2015. Insomnia disorder. Nat Rev Dis Primers 1, 15026.

Moskow, J.J., Bullrich, F., Huebner, K., Daar, I.O., Buchberg, A.M., 1995. Meis1, a PBX1-related homeobox gene involved in myeloid leukemia in BXH-2 mice. Mol Cell Biol 15, 5434–5443.

Nagel, M., Jansen, P.R., Stringer, S., Watanabe, K., de Leeuw, C.A., Bryois, J., Savage, J.E., Hammerschlag, A.R., Skene, N.G., Munoz-Manchado, A.B., andMe Research, T., White, T., Tiemeier, H., Linnarsson, S., Hjerling-Leffler, J., Polderman, T.J.C., Sullivan, P.F., van der Sluis, S., Posthuma, D., 2018. Meta-analysis of genome-wide association studies for neuroticism in 449,484 individuals identifies novel genetic loci and pathways. Nat Genet 50, 920–927.

Pahl, M.C., Doege, C.A., Hodge, K.M., Littleton, S.H., Leonard, M.E., Lu, S., Rausch, R., Pippin, J.A., De Rosa, M.C., Basak, A., Bradfield, J.P., Hammond, R.K., Boehm, K., Berkowitz, R.I., Lasconi, C., Su, C., Chesi, A., Johnson, M.E., Wells, A.D., Voight, B.F., Leibel, R.L., Cousminer, D.L., Grant, S.F.A., 2021. Cis-regulatory architecture of human ESC-derived hypothalamic neuron differentiation aids in variant-to-gene mapping of relevant complex traits. Nat Commun 12, 6749.

Palermo, J., Chesi, A., Zimmerman, A., Sonti, S., Lasconi, C., Brown, E.B., Pippin, J.A., Wells, A.D., Doldur-Balli, F., Mazzotti, D.R., Pack, A.I., Gehrman, P.R., Grant, S.F.A., Keene, A.C., 2021. Variant-to-gene-mapping followed by cross-species genetic screening identifies GPI-anchor biosynthesis as novel regulator of sleep. bioRxiv, 2021.2012.2019.472248.

Parsons, M.J., Lester, K.J., Barclay, N.L., Archer, S.N., Nolan, P.M., Eley, T.C., Gregory, A.M., 2014. Polymorphisms in the circadian expressed genes PER3 and ARNTL2 are associated with diurnal preference and GNbeta3 with sleep measures. J Sleep Res 23, 595–604.

Prober, D.A., Rihel, J., Onah, A.A., Sung, R.J., Schier, A.F., 2006. Hypocretin/orexin overexpression induces an insomnia-like phenotype in zebrafish. J Neurosci 26, 13400–13410.

Rand, K.N., Ho, T., Qu, W., Mitchell, S.M., White, R., Clark, S.J., Molloy, P.L., 2005. Headloop suppression PCR and its application to selective amplification of methylated DNA sequences. Nucleic Acids Res 33, e127.

Reichert, S., Pavon Arocas, O., Rihel, J., 2019. The Neuropeptide Galanin Is Required for Homeostatic Rebound Sleep following Increased Neuronal Activity. Neuron 104, 370–384 e375.

Rihel, J., Prober, D.A., Schier, A.F., 2010. Monitoring sleep and arousal in zebrafish. Methods Cell Biol 100, 281–294.

Ritchey, E.R., Bongini, R.E., Code, K.A., Zelinka, C., Petersen-Jones, S., Fischer, A.J., 2010. The pattern of expression of guanine nucleotide-binding protein beta3 in the retina is conserved across vertebrate species. Neuroscience 169, 1376–1391.

Salminen, A.V., Garrett, L., Schormair, B., Rozman, J., Giesert, F., Niedermeier, K.M., Becker, L., Rathkolb, B., Racz, I., German Mouse Clinic, C., Klingenspor, M., Klopstock, T., Wolf, E., Zimmer, A., Gailus-Durner, V., Torres, M., Fuchs, H., Hrabe de Angelis, M., Wurst, W., Holter, S.M., Winkelmann, J., 2017. Meis1: effects on motor phenotypes and the sensorimotor system in mice. Dis Model Mech 10, 981–991.

Schulte, E.C., Kousi, M., Tan, P.L., Tilch, E., Knauf, F., Lichtner, P., Trenkwalder, C., Hogl, B., Frauscher, B., Berger, K., Fietze, I., Hornyak, M., Oertel, W.H., Bachmann, C.G., Zimprich, A., Peters, A., Gieger, C., Meitinger, T., Muller-Myhsok, B., Katsanis, N., Winkelmann, J., 2014. Targeted resequencing and systematic in vivo functional testing identifies rare variants in MEIS1 as significant contributors to restless legs syndrome. Am J Hum Genet 95, 85–95.

Singh, C., Oikonomou, G., Prober, D.A., 2015. Norepinephrine is required to promote wakefulness and for hypocretin-induced arousal in zebrafish. Elife 4, e07000.

Smemo, S., Tena, J.J., Kim, K.H., Gamazon, E.R., Sakabe, N.J., Gomez-Marin, C., Aneas, I., Credidio, F.L., Sobreira, D.R., Wasserman, N.F., Lee, J.H., Puviindran, V., Tam, D., Shen, M., Son, J.E., Vakili, N.A., Sung, H.K., Naranjo, S., Acemel, R.D., Manzanares, M., Nagy, A., Cox, N.J., Hui, C.C., Gomez-Skarmeta, J.L., Nobrega, M.A., 2014. Obesity-associated variants within FTO form long-range functional connections with IRX3. Nature 507, 371–375.

Spieler, D., Kaffe, M., Knauf, F., Bessa, J., Tena, J.J., Giesert, F., Schormair, B., Tilch, E., Lee, H., Horsch, M., Czamara, D., Karbalai, N., von Toerne, C., Waldenberger, M., Gieger, C., Lichtner, P., Claussnitzer, M., Naumann, R., Muller-Myhsok, B., Torres, M., Garrett, L., Rozman, J., Klingenspor, M., Gailus-Durner, V., Fuchs, H., Hrabe de Angelis, M., Beckers, J., Holter, S.M., Meitinger, T., Hauck, S.M., Laumen, H., Wurst, W., Casares, F., Gomez-Skarmeta, J.L., Winkelmann, J., 2014. Restless legs syndrome-associated intronic common variant in Meis1 alters enhancer function in the developing telencephalon. Genome Res 24, 592–603.

Stranges, S., Tigbe, W., Gomez-Olive, F.X., Thorogood, M., Kandala, N.B., 2012. Sleep problems: an emerging global epidemic? Findings from the INDEPTH WHO-SAGE study among more than 40,000 older adults from 8 countries across Africa and Asia. Sleep 35, 1173–1181.

Su, C., Argenziano, M., Lu, S., Pippin, J.A., Pahl, M.C., Leonard, M.E., Cousminer, D.L., Johnson, M.E., Lasconi, C., Wells, A.D., Chesi, A., Grant, S.F.A., 2021. 3D promoter architecture re-organization during iPSC-derived neuronal cell differentiation implicates target genes for neurodevelopmental disorders. Prog Neurobiol 201, 102000.

Su, C., Johnson, M.E., Torres, A., Thomas, R.M., Manduchi, E., Sharma, P., Mehra, P., Le Coz, C., Leonard, M.E., Lu, S., Hodge, K.M., Chesi, A., Pippin, J., Romberg, N., Grant, S.F.A., Wells, A.D., 2020. Mapping effector genes at lupus GWAS loci using promoter Capture-C in follicular helper T cells. Nat Commun 11, 3294.

Sultmann, H., Sato, A., Murray, B.W., Takezaki, N., Geisler, R., Rauch, G.J., Klein, J., 2000. Conservation of Mhc class III region synteny between zebrafish and human as determined by radiation hybrid mapping. J Immunol 165, 6984–6993.

Tam, V., Patel, N., Turcotte, M., Bosse, Y., Pare, G., Meyre, D., 2019. Benefits and limitations of genome-wide association studies. Nat Rev Genet 20, 467–484.

Thireau, J., Farah, C., Molinari, N., Bouilloux, F., Torreilles, L., Winkelmann, J., Scholz, S., Richard, S., Dauvilliers, Y., Marmigere, F., 2017. MEIS1 variant as a determinant of autonomic imbalance in Restless Legs Syndrome. Sci Rep 7, 46620.

Tran, S., Prober, D.A., 2022. Validation of Candidate Sleep Disorder Risk Genes Using Zebrafish. Front Mol Neurosci 15, 873520.

Vincent, A., Audo, I., Tavares, E., Maynes, J.T., Tumber, A., Wright, T., Li, S., Michiels, C., Consortium, G.N.B., Condroyer, C., MacDonald, H., Verdet, R., Sahel, J.A., Hamel, C.P., Zeitz, C., Heon, E., 2016. Biallelic Mutations in GNB3 Cause a Unique Form of Autosomal-Recessive Congenital Stationary Night Blindness. Am J Hum Genet 98, 1011–1019.

Wallace, M.L., Stone, K., Smagula, S.F., Hall, M.H., Simsek, B., Kado, D.M., Redline, S., Vo, T.N., Buysse, D.J., Osteoporotic Fractures in Men Study Research, G., 2018. Which Sleep Health Characteristics Predict All-Cause Mortality in Older Men? An Application of Flexible Multivariable Approaches. Sleep 41.

Wang, J., Pol, S.U., Haberman, A.K., Wang, C., O’Bara, M.A., Sim, F.J., 2014. Transcription factor induction of human oligodendrocyte progenitor fate and differentiation. Proc Natl Acad Sci U S A 111, E2885–2894.

Watanabe, K., Jansen, P.R., Savage, J.E., Nandakumar, P., Wang, X., andMe Research, T., Hinds, D.A., Gelernter, J., Levey, D.F., Polimanti, R., Stein, M.B., Van Someren, E.J.W., Smit, A.B., Posthuma, D., 2022. Genome-wide meta-analysis of insomnia prioritizes genes associated with metabolic and psychiatric pathways. Nat Genet 54, 1125–1132.

Zhang, G., Neubert, T.A., Jordan, B.A., 2012. RNA binding proteins accumulate at the postsynaptic density with synaptic activity. J Neurosci 32, 599–609.

Zhdanova, I.V., Wang, S.Y., Leclair, O.U., Danilova, N.P., 2001. Melatonin promotes sleep-like state in zebrafish. Brain Res 903, 263–268.

