## Supplementary Figures for "Zebrafish screen of high-confidence effector genes at insomnia GWAS loci implicates conserved regulators of sleep-wake behaviors"

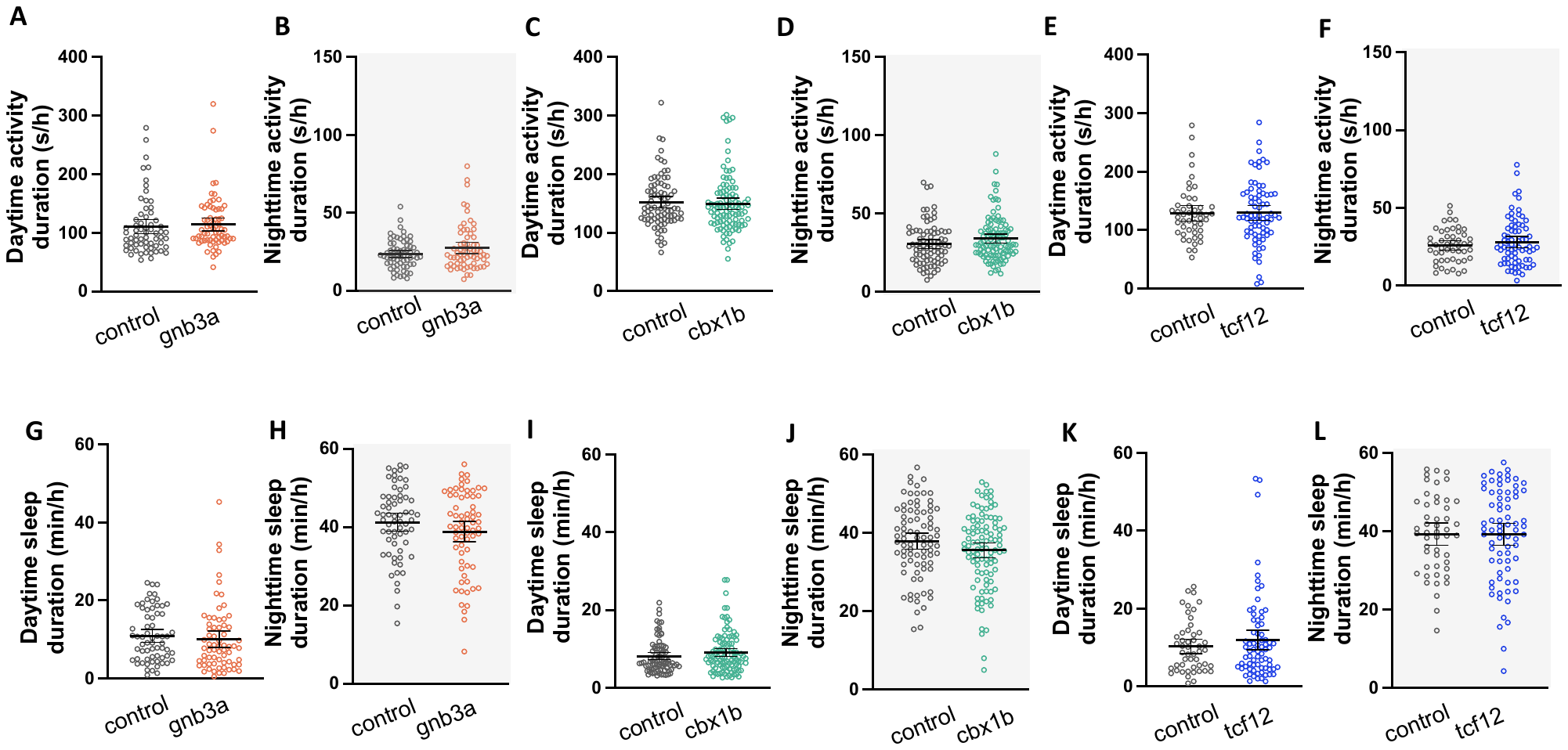

**Supplementary Fig. 1. Sleep and activity duration is not changed in *gnb3a*, *cbx1b*, and *tcf12* mutants.** Mean ± 95% CI for day (**A**) and night (**B**) activity duration in *gnb3a* mutants (n = 64 neg. control, 67 *gnb3a*). Mean ± 95% CI for day (**C**) and night (**D**) activity duration in *cbx1b* mutants (n = 83 neg. control, 102 *cbx1b*). Mean ± 95% CI for day (**E**) and night (**F**) activity duration in *tcf12* mutants (n = 49 neg. control, 73 *tcf12*). Mean ± 95% CI for day (**G**) and night (**H**) sleep duration in *gnb3a* mutants (n = 64 neg. control, 67 *gnb3a*). Mean ± 95% CI for day (**I**) and night (**J**) sleep duration in *cbx1b* mutants (n = 83 neg. control, 102 *cbx1b*). Mean ± 95% CI for day (**K**) and night (**L**) sleep duration in *tcf12* mutants (n = 49 neg. control, 73 *tcf12*).

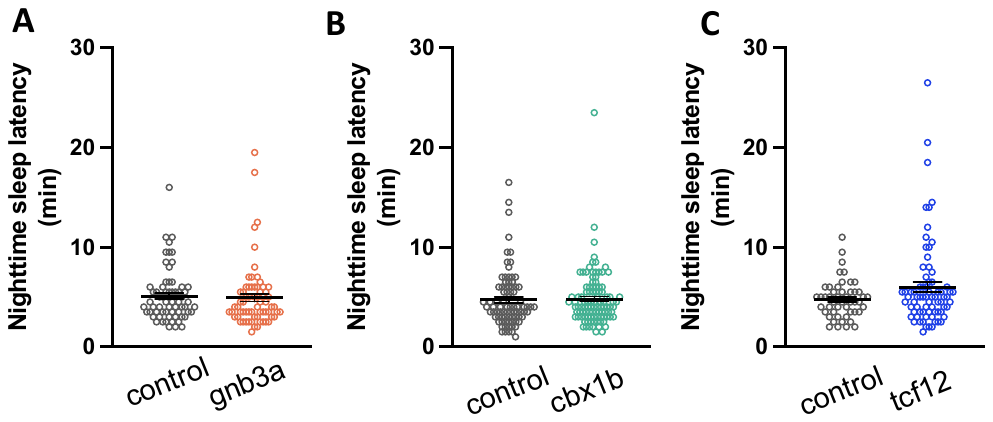

**Supplementary Fig. 2. Latency to sleep onset is not disturbed by mutations in *gnb3a*, *cbx1b*, and *tcf12.*** Mean ± 95% CI for nighttime sleep latency in (A) *gnb3a*, (B) *cbx1b*, and (C) *tcf12*. (n = 64 neg. control, 67 *gnb3a*), (n = 83 neg. control, 102 *cbx1b*)*,* (n = 49 neg. control, 73 *tcf12*).

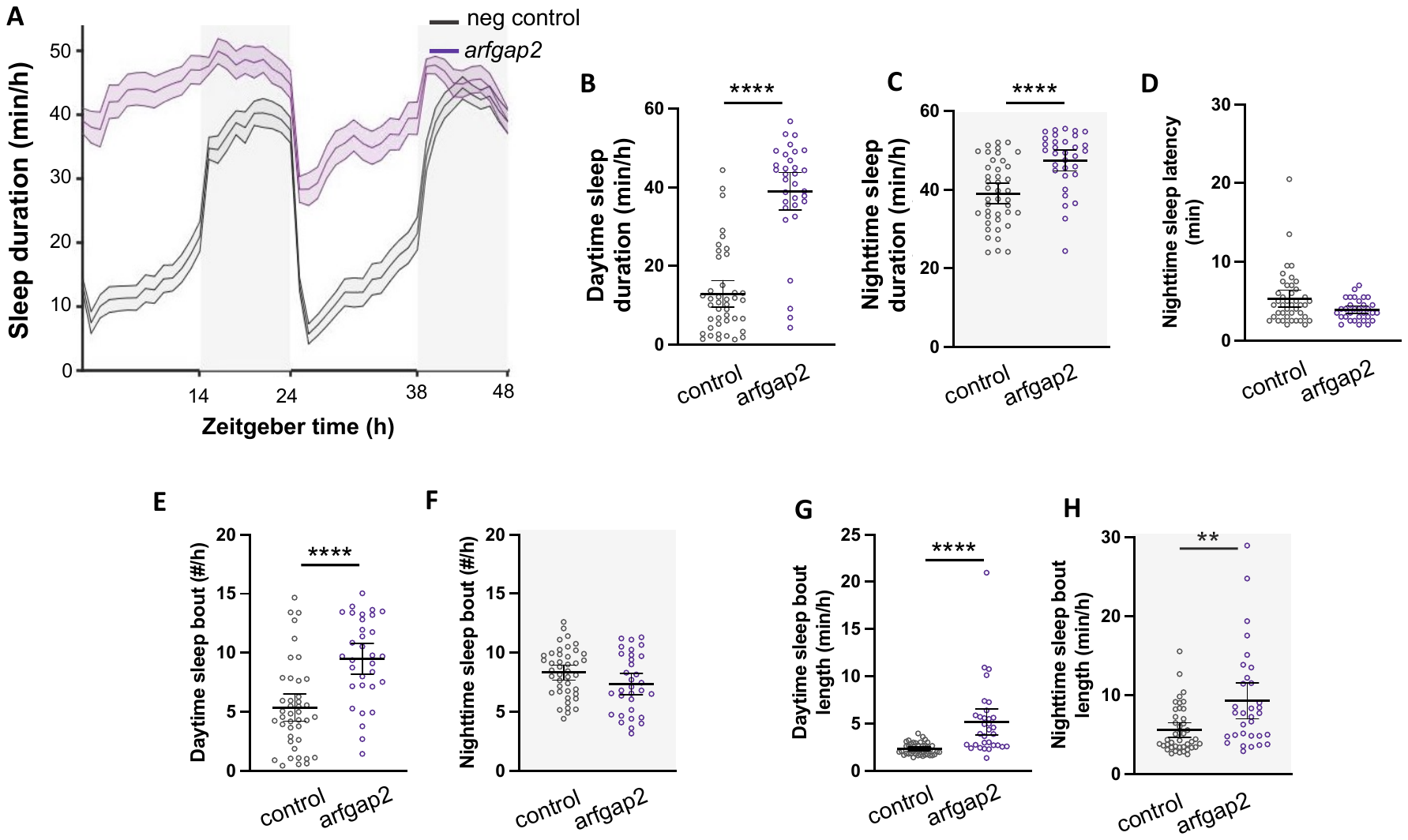

**Supplementary Fig. 3. Mutation of *arfgap2* impairs movement resulting in long bouts of immobility. A.** Rest-activity graph for *arfgap2* mutants represented as mean (line) ± SEM (shaded) for activity duration (min/h). Shaded regions represent lights-off period. Mean ± 95% CI for daytime (**B**) and nighttime (**C**) sleep duration. **D.** Mean ± 95% CI for nighttime sleep latency. Mean ± 95% CI for daytime (**E**) and nighttime (**F**) sleep bout number. Mean ± 95% CI for daytime (**G**) and nighttime (**H**) sleep bout length. n = 42 control, 32 arfgap2. Significance determined by Wilcoxon rank sum test followed by Hochberg step-up procedure for multiple comparisons (see **Methods**). *p<0.05, **p<0.01, ***p<0.001, ****p<0.0001.

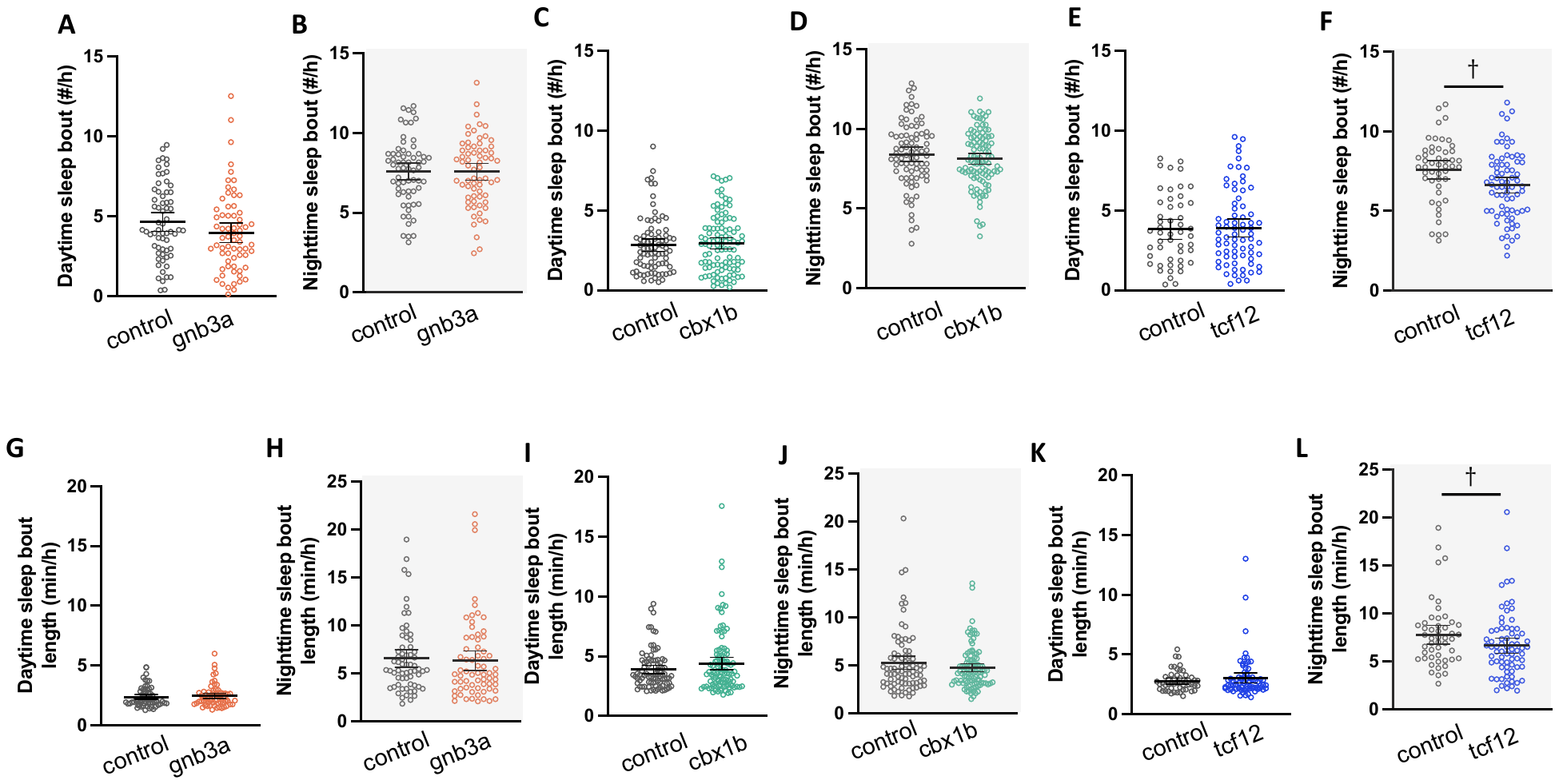

**Supplementary Fig. 4. Sleep characteristics in *gnb3a*, *cbx1b*, and *tcf12* mutants.** Mean ± 95% CI for day (**A**) and night (**B**) sleep bout number in *gnb3a* mutants (n = 64 neg. control, 67 *gnb3a*). Mean ± 95% CI for day (**C**) and night (**D**) sleep bout number in *cbx1b* mutants (n = 83 neg. control, 102 *cbx1b*). Mean ± 95% CI for day (**E**) and night (**F**) sleep bout number in *tcf12* mutants (n = 49 neg. control, 73 *tcf12*). Mean ± 95% CI for day (**G**) and night (**H**) sleep bout length in *gnb3a* mutants (n = 64 neg. control, 67 *gnb3a*). Mean ± 95% CI for day (**I**) and night (**J**) sleep bout length in *cbx1b* mutants (n = 83 neg. control, 102 *cbx1b*). Mean ± 95% CI for day (**K**) and night (**L**) activity sleep bout length in *tcf12* mutants (n = 49 neg. control, 73 *tcf12*). † indicates significant (p < 0.05) before multiple comparisons.

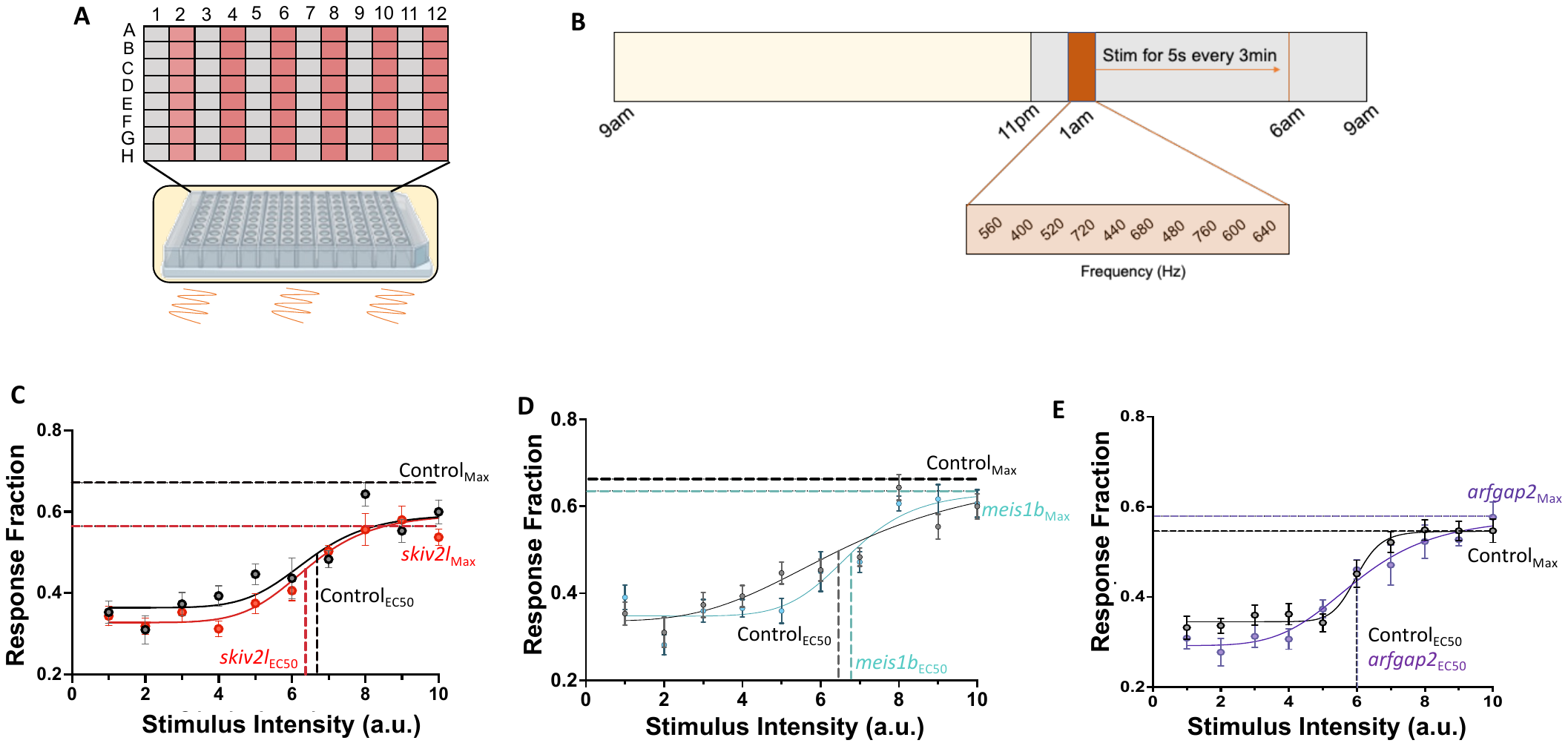

**Supplementary Fig. 5. Arousal threshold is not significantly altered in *skiv2l*, *meis1b*, and *arfgap2* mutants. A.** Cartoon depiction of 96 well plate organization. Different genotypes are placed into alternating columns. Plate is placed on top of a tray within a sound-attenuating chamber (ViewPoint Lifesciences), and vibration stimulus is applied to the bottom of the plate. **B.** Schematic depicting timeline of arousal threshold assay. Animals are kept on a 14:10h light dark cycle with lights off at 11pm. Vibration stimulation occurs for 5s from 1-6am, alternating through 10 pseuodrandomly assigned frequencies every 3 minutes (5sec on: 2:55min off) for 10 total trials at each frequency. **C-E**. Stimulus response curves for (**C**) *skiv2l* (n = 30 control, 32 *skiv2l*), (**D**) *meis1b* (n = 30 control, 32 *meis1b*), and (**E**) *arfgap2* (n = 47 control, 48 *arfgap2*) mutants. Stimulus response curves show EC_50_ (stimulus intensity needed to reach the half-maximal response) and Max response (highest response for the genotype). Error bars represent SEM for average response fraction across all ten trials.
